# Rapid speciation and karyotype evolution in Orthoptera

**DOI:** 10.1101/2021.07.06.451348

**Authors:** Octavio M. Palacios-Gimenez

## Abstract

To test the hypothesis that high speciation rate in groups is coupled with high rate of karyotype evolution but also that younger groups having a higher rate of karyotypic diversity, I estimated rates of speciation and rates of karyotype evolution in 1,177 species belonging to 26 families in the insect order Orthoptera. Rates of karyotype evolution were estimated using the diploid number and the number of chromosome arms (fundamental number) from published karyotypes of Orthoptera. Rates of speciation were quantified considering the number of species examined karyotypically in each family, the most recent common ancestor of each family and the information about extinction rate. The rate of speciation was strongly correlated with rate of karyotype evolution and the average rates of speciation was nearly ~177 times higher than the background rate estimated for Orthoptera based on acoustic communication using phylogenomic data, as well as 8.4 and 35.6 times higher than the estimated speciation rate in vertebrates and bivalve mollusks respectively, indicating that Orthoptera has evolved very fast at chromosomal level. The findings supported the hypothesis of a high speciation rate in lineages with high rate of chromosomal evolution but there were not evidences that younger groups tended to have higher rate of karyotypic diversity. Furthermore, rates of karyotype evolution most closely fitted the punctuational evolutionary model indicating the existence of long periods of stasis of karyotype change with most karyotype change occurring quickly over short evolutionary times. I discussed genetic drift, divergent selection and meiotic drive as potential biological mechanisms to explain karyotype evolution allowing or impeding for the fixation of chromosomal rearrangements and in turn speciation in orthopterans lineages.

## Introduction

The link between karyotype evolution and speciation have long been a topic of heated debate among evolutionary biologists (White 1978; Sites and Moritz 1987; Coyne and Allen Orr 1998; Hoffmann and Rieseberg 2008). Earliest hypothesis pointed out that small populations are essential for quick speciation (Wright 1931, 1940; Mayr 1970; Coyne and Allen Orr 1998). Under this view, a new species might arise when a small subset of individuals isolated from a large population establish a new colony (bottleneck or founder effect) or small demes maintained by social structuring and ecology (Coyne and Allen Orr 1998; Hoffmann and Rieseberg 2008). It is also widespread acceptance of the hypothesis that small population are critical for karyotypic evolution suggesting that speciation and karyotypic change may likely be associated (White 1978; Coyne and Allen Orr 1998; Hoffmann and Rieseberg 2008). It is not easy to obtain direct evidence of the relationship between population size and rate of evolution because estimates of population size is restricted only to a few number of taxa across the Tree of Life. However, it is possible to examine quantitatively the relationship between karyotype evolution and speciation in taxa with well-described karyotypic information. Rates of karyotypic change and speciation might be associated if both factors depend on the occurrence of small population.

The link between karyotypic change and speciation was qualitatively examined in groups of organisms like mammals (Bush et al. 1977), birds (Tegelström et al. 1983), fishes (Yoshida and Kitano 2021) and bivalve mollusks (Stanley 1975), that has led to a heated debate (King 1985; Sites and Moritz 1987). Opinions differ as to the relationship between karyotypic changes and speciation, these changes being considered by some as causative of reproductive isolation and in turn speciation (White 1978; King 1995), and by others as an incidental by-products of speciation processes (Coyne 1984) devoid of any effect on speciation (Futuyma and Mayer 1980; Charlesworth et al. 1982).

Many models of karyotypic changes have been suggested over the past 70 years (Rieseberg 2001). The canalization model (Bickham and Baker 1979) suggested that karyotypic change diversification in younger groups of vertebrates (e.g. rodents) is higher than in older groups of vertebrates (e.g. bats and turtles). This model, however, has been widely criticized because is difficult to test and when prediction were made, empirical evidences do not support the model (see King 1985; Sites and Moritz 1987). The most widely accepted models of karyotypic changes concern the effect of individual chromosomal rearrangements that decrease the fertility of the heterozygotes by either disrupting the segregation pattern in meiosis or reducing the recombination rate in rearranged areas of the genome. Some of the most prominent models within this category differ in both geographic context and in the manner in which new rearrangements are spread once fixed. The stasipatric model (White 1978) assumes that a strongly underdominant chromosomal rearrangement arises and becomes fixed in a population that is within the range of the progenitor. The chain or cascade model (White 1978) alleges that reproductive isolation result from the accumulation of chromosomal rearrangements that are individually underdominant. The chromosomal transilience model (Templeton 1981) states that a strongly underdominant chromosomal rearrangement might be fixed by drift and endogamy in an isolated population. Substantial isolating barrier with respect to the ancestral karyotype might complete speciation. The quantum speciation model (Grant 1981) proposes that chromosomal rearrangements become fixed very quickly in a peripheral founder population by drift and endogamy promoting reproductive isolation. This model is similar to the chromosomal transilience model except that the new gene arrangement resulting from the karyotypic change are thought to be adaptive. The monobraquial fusion model (Baker and Bickham 1986) alleges that different centric fusions become independently fixed in isolated subpopulation; the fusions might then cause little or no loss of fertility on heterozygote individuals. Hybrids between the two subpopulation, however, would be intersterile because different combination of chromosomal arms had been fused in the two subpopulation. Finally, the recombinational model (Lewis 1966) states that hybridization between chromosomal divergent population leads to chromosomal breakages and to the sorting of preexisting rearrangements that differentiate from the parental species. The tenets against these models in speciation are: 1) the observation that most chromosomal rearrangements have little effect on fertility (Dobzhansky 1933; Sites and Moritz 1987; Coyne et al. 1993), 2) theoretical difficulties related to fixing chromosomal rearrangements that are strongly underdominant (i.e. reduce the fitness of heterozygotes) (Walsh 1982; Lande 1985), 3) the supposed ineffectiveness of chromosomal rearrangements to impede gene flow (Barton 1979; Futuyma and Mayer 1980; Spirito, F. 2000), and 4) the widespread acceptance of the idea that premating and/or ecological barriers predate chromosomal rearrangements in the speciation process and thus may likely cause speciation (Coyne et al. 1993; Schluter, D. 1998; Schemske 2000). Much more work is thus needed to test the generalities of these chromosome-based speciation models.

Here I compiled information on diploid number and number of chromosomal arms in 1,177 species of the insect order Orthoptera to investigate two central goals: rates of karyotypic change and speciation in Orthoptera. Rates of karyotypic change was strongly correlated with rate of karyotype evolution. Both rates varied largely in these animals being an order of magnitude in most orthopterans higher than vertebrates and bivalve mollusks. Therefore, it was essential to estimate both rates in a wide variety of orthopterans. The result of such estimates reported bellow were discussed in the light of genetic drift, divergent selection and meiotic drive as potential biological mechanisms to explain karyotype evolution and in turn speciation in orthopterans lineages.

## Methods

### Rates of karyotype evolution

I compiled information on diploid number (2n) and number of chromosome arms (fundamental numbers, FN) from males in 1,177 orthopteran species belonging to 25 families (Fig. 1), based on published data from the literature (Supplementary Table S1). The compiled data concerned families of long-horned grasshoppers (suborder Ensifera) and short-horned grasshoppers (suborder Caelifera), and therefore, covered the Orthoptera Tree of Life. Only diploid species were considered. Polymorphic species for diploid numbers were taking into account. B chromosomes were not considered. Utilising this information, I estimated the rates of karyotype evolution (*r’*) within a major taxonomic group (i.e., family) following the equation proposed elsewhere (Bush et al. 1977):

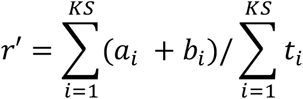

in which *KS* is the number of species examined karyotypically within a family, and *t* is the divergence time of the most recent common ancestor of a family in the fossil record (Song et al. 2015, 2020) in millions of years ago (Myr). The symbol *a* was defined as (*c- d)/KS*, in which *c* stands for the highest diploid number and *d* for the lowest diploid number within the family. Likewise, *b* was defined as (*e- f*)/*KS*, in which *e* stands for the highest FN and *f* for the lowest FN. Thus, the rate quantifies karyotypic changes/per lineages per million years. The method deals with chromosomal rearrangements like fusions/fissions and pericentromeric inversions that alter chromosome number and the FN, respectively. However, it doesn’t deal with most micro-chromosomal rearrangements (e.g. paracentric inversions) and chromosomal reciprocal translocations that might be critical for karyotype evolution, reproductive isolation and in turn speciation. The latter mutations are usually detected in vertebrates through chromosome painting and chromosome banding techniques, but unfortunately, this information is not available in the orthopterans cytogenetic literature because it is difficult to recognize through classical cytogenetic methods, and therefore, these mutations were not taking into account.

**Figure 1:**
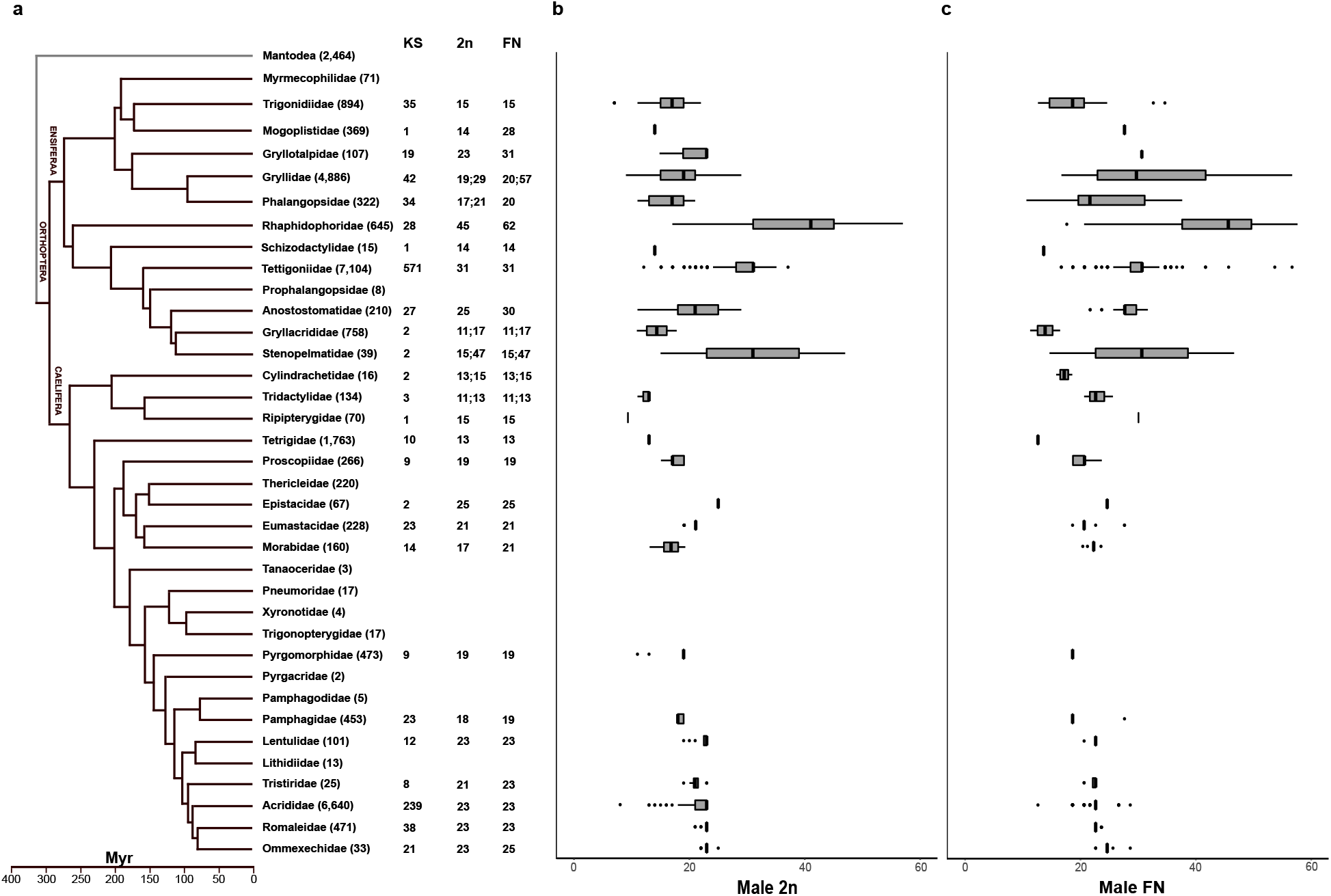
Karyotype and fundamental number distributions across the orthoptera phylogeny. **a** Dated phylogeny of Orthoptera based on phylogenomic data (Song et al. 2015, 2020). Each terminal represents a monophyletic family and the number in parenthesis next to the family name indicates the number of validly described species within the family. The number of karyotyped species (KS), the most common diploid number (2n) and most common fundamental number (FN) in males are showed for the 25 analysed families. **b** and **c** Boxplots showing the distribution of both the diploid number (2n) and the fundamental number (FN) in males across each family analysed. Myr= millions of years ago.

### Rates of speciation

Speciation rate (*S*) within family of Orthoptera was estimated by using *S*= *R* + *E* (Stanley 1975). Within a family, *R* stands for the net rate at which a new species have arisen, and *E* stands for the average rate at which species have become extinct over a defined period of time. The mean value of the *R* was estimated for each family of Orthoptera following the equation described elsewhere (Bush et al. 1977):

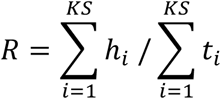

in which *KS* is the number of species examined karyotypically within a family, and *h* stands for the number of speciation events per lineage. Therefore, the *h* values were summed and divided by the summed time range (*t*) for the family. The *h* values were estimated by using the following equation (Bush et al. 1977):

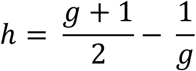

in which *g* stands for the number of existing species per family obtained from the literature (Song et al. 2015; Cigliano et al. 2021).

To estimate the average extinction rate (*E*) for species within the family, I first looked back in geological time to find strata in which 50% of the family fossils belong to extant species. The time was then designed *D*/2, in which *D* was defined to an estimate of the species duration for species in that family. The mean extinction rate (*E*) was then defined to be 1/D (Stanley 1975; Bush et al. 1977). Divergence time were obtained from the fossil record available in the literature (Song et al. 2015, 2020). Finally, I got an estimate of *S* (referred to as the corrected speciation rate) for each family by adding the *E* and *R* values. All of the statistical analysis were ran in R version 3.5.1 (R Core Team 2018).

## Results

The 2n across males of Orthoptera varied from 7 in *Eunemobius carolinus carolinus* (Trigoniidae) to 57 in *Diestrammema tachycines asynamorus* (Rhaphidophoridae), while the NF varied from 11 in *Strinatia brevipenis* (Phalangopsidae) and *Gryllacris signifera* (Gryllacrididae) to 64 in *Dolichopoda schiavazzii* (Rhaphidophoridae) (Fig. 1, Supplementary material 1). The karyotype diversity (Shannon index, *H’*) estimated from the FN varied from 0 (Mogoplistidae, Schizodactylidae, Ripipterygidae) to 6.341 (Tettigoniidae). The karyotypic changes/per lineages per million years (*r’*) estimated for all of the families analysed appear in Table 1. A linear regression indicated that *r’* is determined by the percentage of *KS* per family (*P*= 0.0107), indicating that in the collected data set this mutation rate is sensitive to sampling bias. Thus, *r’* in families with lower *KS* were likely underestimated. The highest *r’* occurred within Tettigoniidae and within the recent-divergent lineages Gryllidae, Phalangopsidae, Acridiade, Romaleidae and Ommexechidae, indicating that orthopterans have evolved very fast at chromosomal level.

**Table 1:** Rates of karyotype evolution in extant family of Orthoptera

| Family | Number of<br>karyotyped species,<br>*KS* | FN Shannon<br>index, *H'* | Average age of<br>the family,<br>Myr | Karyotypic changes/<br>lineage per Myr, *r'* |
| --- | --- | --- | --- | --- |
| Trigonidiidae | 35 | 3.525 | 175 | 0.322 |
| Mogoplistidae* | 1 | 0 | 200 | 0 |
| Gryllotalpidae | 19 | 2.944 | 98 | 0.526 |
| Gryllidae | 43 | 3.696 | 98 | 0.871 |
| Phalangopsidae | 34 | 3.488 | 98 | 0.595 |
| Schizodactylidae* | 1 | 0 | 215 | 0 |
| Tettigoniidae | 571 | 6.341 | 115 | 0.816 |
| Rhaphidophoridae | 28 | 3.297 | 215 | 0.443 |
| Anostomatidae | 27 | 3.292 | 125 | 0.478 |
| Gryllacrididae | 2 | 0.670 | 150 | 0.153 |
| Stenopelmatidae | 2 | 0.553 | 125 | 0.632 |
| Cylindrachetidae | 2 | 0.693 | 202 | 0.103 |
| Tridactylidae | 3 | 1.095 | 320 | 0.088 |
| Ripterygidae* | 1 | 0 | 153 | 0 |
| Tetrigidae | 10 | 2.301 | 263 | 0.089 |
| Proscopiidae | 9 | 2.193 | 167 | 0.234 |
| Epistacidae | 2 | 0.693 | 167 | 0.149 |
| Eumastacidae | 23 | 3.131 | 162 | 0.292 |
| Morabidae | 14 | 2.633 | 162 | 0.259 |
| Pyrgomorphidae | 9 | 2.197 | 147 | 0.235 |
| Pamphagidae | 23 | 3.131 | 125 | 0.363 |
| Lentulidae | 12 | 2.485 | 80 | 0.533 |
| Tristiridae | 8 | 2.079 | 98 | 0.418 |
| Acrididae | 238 | 5.475 | 80 | 0.648 |
| Romaleidae | 38 | 3.637 | 80 | 0.573 |
| Ommexechidae | 21 | 3.045 | 77 | 0.673 |
| **Average** |  |  |  | **0.365** |

\*underestimated rate owing to the small sampling.

My estimates of the speciation rate (*S*) within the analysed families appear in Table 2. The lowest net of speciation (*R*) occurred in Tridactylidae and Cylindrachetidae and the highest one in Acrididae, Gryllidae and Tettigoniidae. Extinction rate (*E*) was lower on average in Orthoptera than in mammals and birds. Consequently, the corrected rates of speciation (*S*) were substantially higher in Orthoptera than in vertebrates. Acrididae, Gryllidae and Tettigoniidae experienced substantially high *S* compared to the other families. To test whether high speciation rate in groups is coupled with high rate of karyotype evolution, I plotted *S* against *r’* for the analysed group of Orthoptera (Fig. 2). A linear regression indicated that *S* is determined by the percentage of *r’* per family (*P*= 0.00596). In other words, families with higher *r’* experienced higher *S*, e.g. Tettigoniidae, Gryllidae, Acrididae.

**Table 2:**
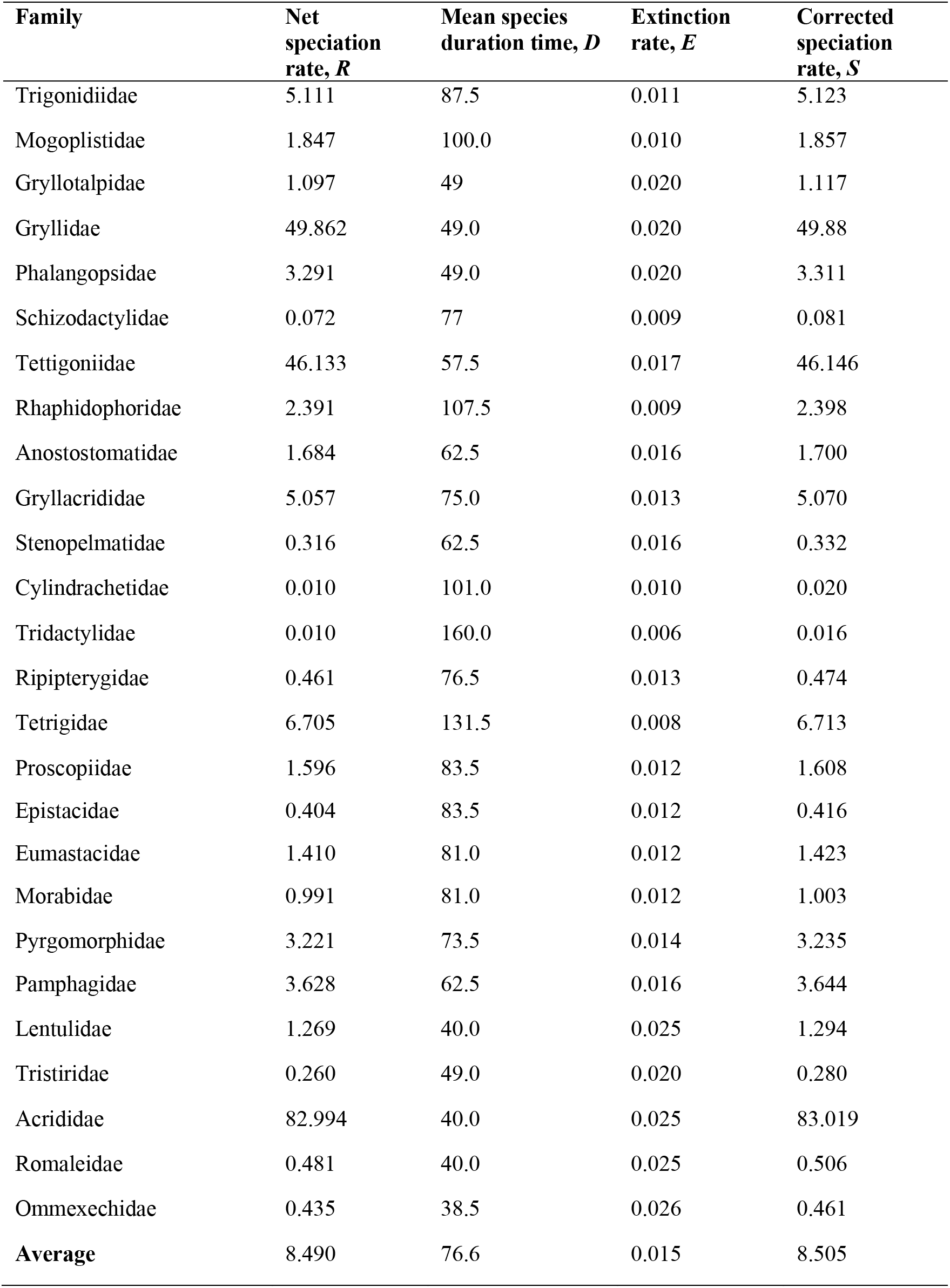
Rates of speciation in extant family of Orthoptera

**Figure 2.**
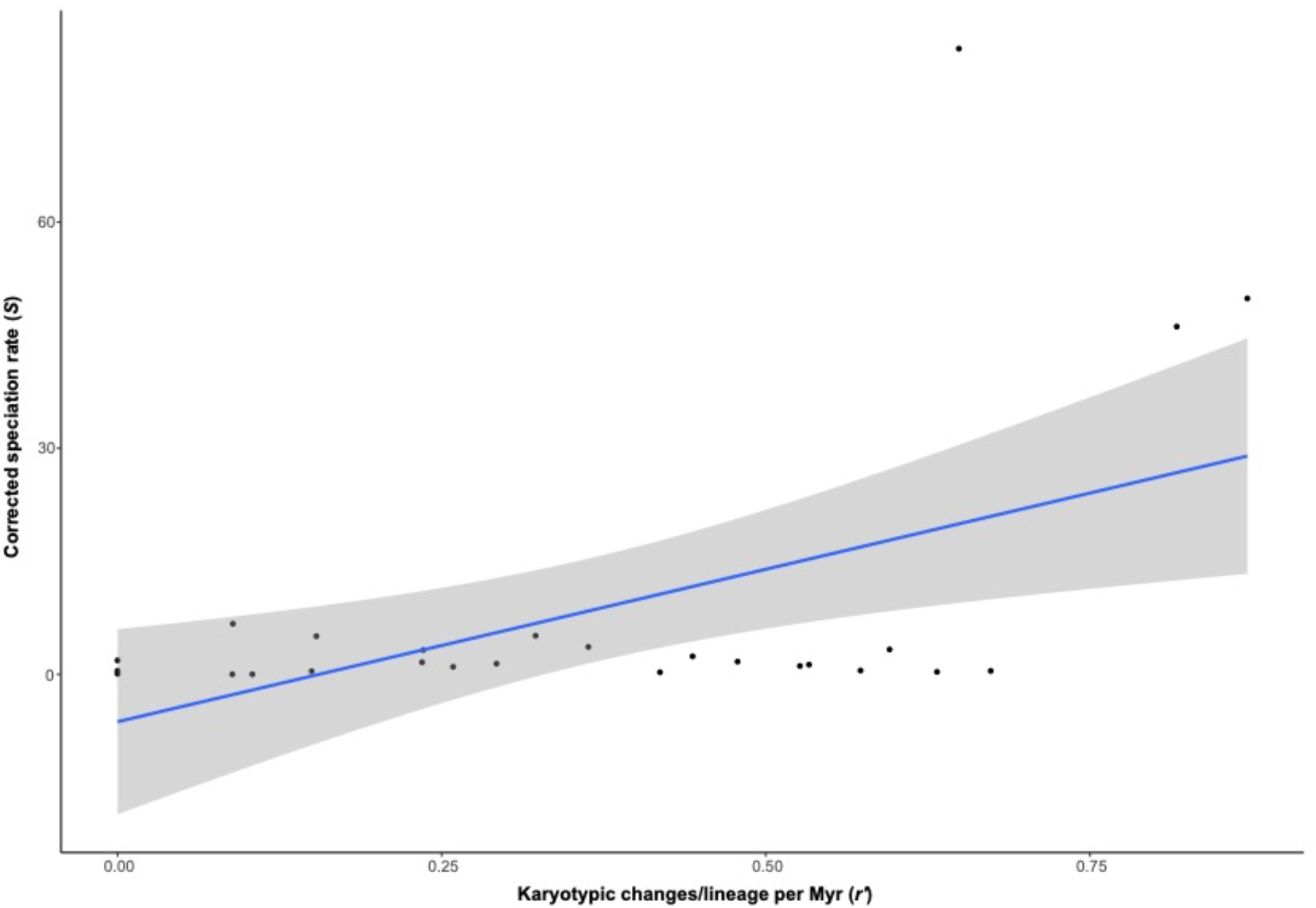
Linear regression showing the relationship between corrected speciation rate (*S*) and rate of karyotypic changes (*r’*) for the 26 families of Orthoptera analysed (*P*= 0.00596). Myr= million years.

## Discussion

The results obtained highlight the existence of a general correlation between karyotypic changing rate and speciation rate in Orthoptera. Rapid karyotypic evolution in Orthoptera (average 0.367 events/lineage per Myr) compared to Polyneoptera insects (average 0.003-0.128) (Sylvester et al. 2020), vertebrates (average 0.166) (Bush et al. 1977; Yoshida and Kitano 2021) and non-passerine birds (average 0.167) (Tegelström et al. 1983) was supported by the high number of species with dissimilar karyotypes. Remarkably, the average karyotypic change reported here was 122 times higher than what has been recently reported in Orthoptera (average 0.003) (Sylvester et al. 2020). This difference is likely owing to the higher number of sampled species in this study, and because ii) the previous report only considered the variation in 2n to estimate the karyotypic changes. It was therefore essential the increasing of the number of sampled species and the compiling of FN to get a better estimation of the karyotypic change in Orthoptera.

Importantly, there were not evidences that younger groups tended to have higher rate of karyotypic changes as alleged by the canalization model (Bickham and Baker 1979). On the contrary, I have observed the reverse pattern alleged by this model: younger group like Gryllidae (divergence time <98 Myr) and the modern grasshoppers like Acrididae, Romaleidae (divergence time <80 Myr) and Ommexechidae (divergence time <77 Myr) had lower rate of karyotypic change than older group like Tettigoniidae (divergence time ~154 Myr). Furthermore, Orthoptera seemed to have limited extinction rate across its Tree of Life. The higher speciation rate observed in Acrididae, Gryllidae and Tettigoniidae can be accepted as a satisfactory measures of the evolutionary success of these groups. Acrididae grasshoppers represent one of most recently diverged lineages within Orthoptera (Song et al. 2020) and seemed to have radiated quickly with comparably higher speciation rate than other orthopterans, with little or no extinction similar to that observed in the oldest Tettigoniidae. Importantly, Acrididae, Gryllidae and Tettigoniidae seems not only to have a comparably high rate of changes in number of chromosomes but also a comparably high number of centric fusions and pericentromeric inversions, indicating that these genomes have high potential for change while the others remained frozen in time, in agreement with the punctuational evolutionary model (Eldredge and Gould 1972). Therefore, centric fusions and pericentromeric inversions played a critical role in karyotype reshuffling as demonstrated by the reduction in male 2n and in male FN variation. Among these, centric fusions tend to be the most common rearrangements. Fissions rate were reported to be lower in Orthoptera (average 0.024) compared to Polyneoptera insects (average 0.063-0.150) (Sylvester et al. 2020) most likely because of the ineffectiveness of chromosomes to form or assemble new centromeric structures essential for the proper chromosome segregation during cell divisions (Barra and Fachinetti 2018).

But why were the centric fusions the most common large-scale chromosomal rearrangements in Orthoptera? In general, Orthoptera has very large (9 Gb on average; (Gregory 2021)) and repeat-enriched genomes mostly because the expansion of repetitive sequences, ranging from 33 to 75% per genome assembly (Palacios-Gimenez et al. 2020a; Ylla et al. 2020). Chromosomal rearrangements might essentially depend on these repeated sequences that are recombination hot spots in groups with high recombination rates such as rodents having large portion of repeats in pericentromeric regions promoting chromosomal rearrangements (Bailey et al. 2004). In Orthoptera, pericentromeric regions are heterochromatic and very enriched of highly-homogeneous satellite DNA sequences (Ruiz-Ruano et al. 2016; Milani et al. 2018; Palacios-Gimenez et al. 2020b) while transposable elements (TEs) are mostly scattered in the euchromatin of chromosomes, as demonstrated by *in situ* hybridization (Montiel et al. 2012; Palacios-Gimenez et al. 2014; Martí et al. 2021). The divergent pattern concerned the location of satellite DNA and TEs might likely be an indication that of karyotypic intrinsic characteristics (e.g. heterochromatin and satDNA localization) modulate chromosomal fusions; however this idea was never tested in Orthoptera. If this hold true, it might be possible that centric fusions dependent on satellite DNA sequences (White 1978; Dover 2002), although the determinant factors allowing or impeding the fixation of karyotypic changes will be natural selection, genetic drift and/or meiotic drive (see below). It might also be possible that repetitive DNA other than satellite DNA (e.g TEs) promotes micro-structural rearrangements (e.g. deletions and insertions), paracentric inversions and chromosomal reciprocal translocations by non-homologous recombination that might be critical for karyotype evolution, reproductive isolation and in turn speciation. These kind of mutations remained undetectable under the methodology used here. High-quality chromosome-scale assemblies with in-depth repeat and gene annotation not yet available for orthopterans are important steps ahead to provide accurate description of these kind of mutations in Orthoptera.

The correlation between the rate of karyotype evolution and speciation in Orthoptera may be promoted by less vagility in some species and consequently by the formation of reproductively isolated demes also in the absence of geographical separation. The brachypterus South American Melanoplinae grasshopper *Dichroplus silveiraguidoi* (male 2n= 8, neo-XY, FN= 13) form but one example of the extreme divergence from the modal 2n and FN in Acrididae grasshoppers, in which the karyotype have been largely rearranged by centric fusions that involved autosome-autosome and autosome-X chromosomes (Saez 1957; Saez and Perez-Mosquera 1977). The flightless Australian Morabidae grasshoppers in which the karyotypes have been largely reshuffled (male 2n from 14 to 19) by chromosomal rearrangements represent an example of extensive karyotype variation in a parapatric distribution pattern, and narrow hybrid zones at their boundaries. (White 1978; Kawakami et al. 2009). Among long-horned grasshoppers, the cricket *Eneoptera surinamensis* (male 2n= 9, neo-X1X2Y, FN= 17) (Palacios-Gimenez et al. 2015) restrict to Southern America and Northern South America (Cigliano et al. 2021), represent an example of the extreme divergence from the modal 2n and FN in Gryllidae (see Fig. 1). To my knowledge, there is not such karyotypes divergence in orthopterans with either great flight capacity or global distribution, apart from the South American grasshopper *Dichroplus pratensis* (restricted to Argentina, Uruguay and Southern Brazil) which is polymorphic and polytypic for a system of centric fusions (Bidau and Martí 2002). Fixation of multiple centric fusions and consequently greater karyotype divergence in the species abovementioned might have been favored by its less vagility. Furthermore, multiple chromosomal mutations might have impeded genic introgression via hybridization between species that differentiated by allopatry in agreement with the model of chromosomal primary allopatric speciation (King 1995). If this hold true, the fixation of the multiple centric fusions might have predated the genetical and morphological differences (King 1995). Cases of species differentiating allopatrically have been documented in reptiles, in which genic introgression in cases of secondary contact with the parental species was impeded by chromosome differences (King 1995).

Though karyotypic changes are not a condition for speciation, it is widespread acceptance that karyotypic change may contribute to the establishment of reproductive isolation favoring the differentiation of incipient species (Faria and Navarro 2010). But how karyotypic changes may become fixed in populations leading to differentiation and in turn speciation? White (White 1978) alleged four factor that acting separately or combined might have influence in the incorporation of a chromosomal variant in populations: genetic drift, meiotic drive, selective advantage of the homokaryotype and inbreeding. The local chromosomal characteristic may diverge from the parental condition by the action of either adaptation or genetic drift. It is widespread acceptance that different environments produce differential selective pressure. Hence, if a karyotypic change confers an adaptative advantage to carriers in a given condition, it might likely spread and eventually become fixed in a population (Kirkpatrick and Barton 2006; Hooper and Price 2015). Furthermore, species with patchy distribution and small demes may favor the chance of fixation of the rearranged chromosomes by genetic drift, although gene flow between population will be the major factor determining the fixation of local adaptive traits (Hooper and Price 2015). Lowly vagile species like the brachypterous *D. silveiraguidoi* tends to have few migrants between populations and, and consequently, a higher probability of fixation of adaptive traits. In addition, if the selective pressure on the rearranged chromosome is very strong it might even become fixed in the presence of gene flow (Coyne and Orr 2004; Kawakami et al. 2009). This may be the case of centric fusions and pericentromeric inversions having pronounced effect on meiotic recombination by protecting coadapted alleles (supergenes) or by distorting the laws of Mendelian inheritance (meiotic drive) that can be beneficial for the carriers in certain conditions (Bidau et al. 2001; Avril et al. 2020). Under this view, a global geographical distribution of families like Tettigoniidae Gryllidae, Acrididae might expose its species to different selective pressures allowing the fixation of different chromosomal rearrangements in different conditions.

However, it is also widespread acceptance that most chromosomal rearrangements are either not adaptative (Hoffmann and Rieseberg 2008; Faria and Navarro 2010) or have little effect on fertility (Sites and Moritz 1987; Coyne et al. 1993). However, for this to happen is essential that populations with different combination of rearranged chromosomes are isolated geographically (Lande 1985). The theoretical prediction is that population are more prone to diverge genetically in allopatry, and under certain conditions, to accumulate chromosomal rearrangements (Faria and Navarro 2010). Populations are prone to differentiate in the peripheral areas of the distribution of the species because they have the chance to colonize and exploit new habitats or niches (Bush et al. 1977). If an underdominant or neutral chromosomal rearrangement arises under this condition, the rearranged chromosome in that population might differentiates and expands if they are geographically isolated concerned to the parental population (Wright 1949; Lande 1985). On the contrary, they will coalesce with the more numerous parental population via hybridization, eventually the new mutation will be lost quickly because the lack of adaptative advantage that might allow its permanence (Hoffmann and Rieseberg 2008). Again, low-vagility species might favor this diversification mechanism because this condition might allowed exclusive variants to become isolated and fixed in different localities, leading to speciation. The highest rates of karyotypic change and speciation found in Acrididae, Gryllidae and Tettigoniidae are perfect to test this ideas. Among these, the South American Melanoplinae (Acrididae) grasshoppers might be the most suitable species because of their disproportionate frequency of rearranged karyotypes derived from centric fusions than other orthopterans (Castillo et al. 2010).

In summary, it was essential to estimate rates across the Orthoptera Tree of Life. The findings showed that karyotype evolution mediated by chromosomal rearrangements strongly correlates with speciation rate, likely indicating that karyotype evolution is not neutral. I further showed that speciation rate is in-order-of-magnitude higher than vertebrates and bivalve mollusks, and nearly ~177 times higher than the estimated background rate of Orthoptera based on acoustic communication using phylogenomic data (Song et al. 2020). Chromosomal rearrangements like centric fusions are very common in Orthoptera. Centric fusions might have substantial and immediate evolutionary impact by reducing genetic diversity in rearranged areas of the genome likely reflecting increased background selection and selection against introgression between diverging small demes or population, following a reduction in recombination rate. I also provided evidences that karyotype evolution in Orthoptera most closely fitted the punctuational model of evolution indicating the existence of long periods of stasis of karyotype change with most karyotype change occurring quickly over short evolutionary times. Finally, the rapid karyotypic changes in Orthoptera suggest that this taxa is a treasure box for studying the implication of chromosomal rearrangements in evolution and speciation, and deserves more attention in this respect.

## Ethics

Not applicable.

## Data accessibility

The datasets generated as part of the study are available as supplementary information.

## Acknowledgements

The manuscript was improved by comments from Diogo C. Cabral-de-Mello. Computations were performed on resources provided by the Swedish National Infrastructure for Computing (SNIC) through the Uppsala Multidisciplinary Center for Advanced Computational Science (UPPMAX).

