## Supplementary material for "Rapid speciation and karyotype evolution in Orthoptera"

**Table S1.** Species utilised to estimate the rate of karyotypic change for families. FN= fundamental number or number of chromosome arms. Unknowing FN were highlighted in grey, and in this cases was assumed that FN is equal to the 2n of the species.

| **Family** | **Species** | **Male 2n** | **FN** | **References** |
|---|---|---|---|---|
| Trigonidiidae | *Allonemobius allardi* | 15 | 15 | (Portugal and Mesa 2007) |
| Trigonidiidae | *Allonemobius griseus griseus* | 15 | 15 | (Portugal and Mesa 2007) |
| Trigonidiidae | *Allonemobius fasciatus* | 15 | 15 | (Portugal and Mesa 2007) |
| Trigonidiidae | *Allonemobius maculatus* | 15 | 15 | (Portugal and Mesa 2007) |
| Trigonidiidae | *Allonemobius tinnulus* | 15 | 15 | (Portugal and Mesa 2007) |
| Trigonidiidae | *Dianemobius chibae* | 15 | 15 | (Portugal and Mesa 2007) |
| Trigonidiidae | *Dianemobius (Dianemobius) csikii* | 17 | 17 | (Portugal and Mesa 2007) |
| Trigonidiidae | *Dianemobius (Dianemobius) fascipes* | 17 | 18 | (Portugal and Mesa 2007) |
| Trigonidiidae | *Dianemobius (Dianemobius) furumagiensis* | 19 | 23 | (Portugal and Mesa 2007) |
| Trigonidiidae | *Dianemobius mikado* | 15 | 25 | (Portugal and Mesa 2007) |
| Trigonidiidae | *Dianemobius (Polionemobius) flavoantennalis* | 17 | 21 | (Portugal and Mesa 2007) |
| Trigonidiidae | *Dianemobius (Polionemobius) taprobanensis* | 15 | 25 | (Portugal and Mesa 2007) |
| Trigonidiidae | *Eunemobius carolinus carolinus* | 7 | 13 | (Portugal and Mesa 2007) |
| Trigonidiidae | *Eunemobius confusus* | 7 | 13 | (Portugal and Mesa 2007) |
| Trigonidiidae | *Eunemobius melodius* | 7 | 13 | (Portugal and Mesa 2007) |
| Trigonidiidae | *Nemobius sylvestris* | 17 | 33 | (Portugal and Mesa 2007) |
| Trigonidiidae | *Neonemobius variegatus* | 19 | 21 | (Portugal and Mesa 2007) |
| Trigonidiidae | *Neonemobius cubensis* | 19 | 19 | (Portugal and Mesa 2007) |
| Trigonidiidae | *Neonemobius eurynotu* | 19 | 19 | (Portugal and Mesa 2007) |
| Trigonidiidae | *Neonemobius palustris* | 19 | 19 | (Portugal and Mesa 2007) |
| Trigonidiidae | *Phoremia circumcincta* | 17 | 19 | (Portugal and Mesa 2007) |
| Trigonidiidae | *Phoremia nigrofasciata* | 21 | 25 | (Portugal and Mesa 2007) |
| Trigonidiidae | *Pictonemobius sp.* | 19 | 19 | (Portugal and Mesa 2007) |
| Trigonidiidae | *Pteronemobius (Pteronemobius) birmanus* | 15 | 15 | (Portugal and Mesa 2007) |

| | | | | |
|---|---|---|---|---|
| Trigonidiidae | *Pteronemobius (Pteronemobius) nigriscens* | 17 | 17 | (Portugal and Mesa 2007) |
| Trigonidiidae | *Pteronemobius (Pteronemobius) nigrofasciatus* | 17 | 20 | (Portugal and Mesa 2007) |
| Trigonidiidae | *Pteronemobius (Pteronemobius) nitidus* | 17 | 17 | (Portugal and Mesa 2007) |
| Trigonidiidae | *Pteronemobius (Pteronemobius) ohmachii* | 11 | 21 | (Portugal and Mesa 2007) |
| Trigonidiidae | *Nemobius yezoensis* | 17 | 19 | (Portugal and Mesa 2007) |
| Trigonidiidae | *Stenonemobius(Ocellonemobius) bicolor* | 19 | 19 | (Portugal and Mesa 2007) |
| Trigonidiidae | *Scottiola sp.* | 15 | 15 | (Portugal and Mesa 2007) |
| Trigonidiidae | *Zucchiella atlantica* | 22 | 35 | (Portugal and Mesa 2007) |
| Trigonidiidae | *Anaxipha pallidula* | 17 | 17 | (White 1973) |
| Trigonidiidae | *Anaxipha pallidula* | 19 | 19 | (White 1973) |
| Trigonidiidae | *Trigonidium cicindeloides* | 11 | 22 | (Handa et al. 1985) |
| Mogoplistidae | *Cycloptiloides americanus* | 14 | 28 | (Palacios-Gimenez and Cabral-de-Mello 2015) |
| Gryllidae | *Acheta desertus* | 21 | 42 | (Nilsson et al. 2009) |
| Gryllidae | *Acheta domesticus* | 21 | 42 | (Nilsson et al. 2009) |
| Gryllidae | *Brachytrupes portentosus* | 13 | 26 | (White 1973) |
| Gryllidae | *Brachytrupes portentosus* | 14 | 28 | (White 1973) |
| Gryllidae | *Brachytrupes portentosus* | 15 | 30 | (White 1973) |
| Gryllidae | *Brachytrupes portentosus* | 16 | 32 | (White 1973) |
| Gryllidae | *Brachytrupes portentosus* | 17 | 34 | (White 1973) |
| Gryllidae | *Gryllus assimilis* | 29 | 57 | (Palacios-Gimenez et al. 2015) |
| Gryllidae | *Gryllus argentinus* | 29 | 57 | (Drets and Stoll 1974) |
| Gryllidae | *Gryllus campestris* | 29 | 57 | (Warchałowska-Sliwa 1980) |
| Gryllidae | *Gryllus bimaculatus* | 29 | 57 | (Warchałowska-Sliwa 1980) |
| Gryllidae | *Gryllus confirmatus* | 19 | 38 | (Handa et al. 1985) |
| Gryllidae | *Gryllus desertus* | 21 | 42 | (Honda 1926) |
| Gryllidae | *Gryllus domesticus* | 21 | 42 | (Honda 1926) |
| Gryllidae | *Gryllus nitratus* | 25 | 50 | (Honda 1926) |
| Gryllidae | *Gryllus melanocephalus* | 21 | 42 | (Handa et al. 1985) |

| | | | | |
|---|---|---|---|---|
| Gryllidae | *Gryllodes berthellus* | 23 | 46 | (Honda 1926) |
| Gryllidae | *Gryllodes sigillatus* | 21 | 42 | (Handa et al. 1985) |
| Gryllidae | *Loxoblemmus animao* | 15 | 30 | (Handa et al. 1985) |
| Gryllidae | *Loxoblemmus deteerus* | 11 | 24 | (Handa et al. 1985) |
| Gryllidae | *Loxoblemmus arientalus* | 13 | 26 | (Hewitt 1979) |
| Gryllidae | *Loxoblemmus orientalus* | 14 | 28 | (Hewitt 1979) |
| Gryllidae | *Loxoblemmus orientalus* | 15 | 30 | (Hewitt 1979) |
| Gryllidae | *Loxoblemmus orientalus* | 16 | 32 | (Hewitt 1979) |
| Gryllidae | *Loxoblemmus orientalus* | 17 | 34 | (Hewitt 1979) |
| Gryllidae | *Loxoblemmus doenitzi* | 11 | 22 | (Honda 1926) |
| Gryllidae | *Loxoblemmus sp.* | 13 | 26 | (Hewitt 1979) |
| Gryllidae | *Stephoblemmus humbertiellus* | 21 | 24 | (Handa et al. 1985) |
| Gryllidae | *Teleogryllus commodus* | 27 | 52 | (Lim et al. 1969) |
| Gryllidae | *Teleogryllus oceanicus* | 27 | 52 | (Lim et al. 1969) |
| Gryllidae | *Turanogryllus jammuensis* | 19 | 38 | (Handa et al. 1985) |
| Gryllidae | *Eneoptera surinamensis* | 9 | 17 | (Palacios-Gimenez et al. 2015) |
| Gryllidae | *Duolandrevus (Duolandrevus) brachypterus* | 19 | 20 | (Gorochov and Warchałowska-Śliwa 2004) |
| Gryllidae | *Duolandrevus (Duolandrevus) coulonianus* | 19 | 20 | (Gorochov and Warchałowska-Śliwa 2004) |
| Gryllidae | *Duolandrevus (Bejorama) modestus sp. n* | 19 | 20 | (Gorochov and Warchałowska-Śliwa 2004) |
| Gryllidae | *Duolandrevus (Bejorama) improvisus sp. n.* | 19 | 20 | (Gorochov and Warchałowska-Śliwa 2004) |
| Gryllidae | *Ectodrelanva paramarginalis* | 21 | 22 | (Gorochov and Warchałowska-Śliwa 2004) |
| Gryllidae | *Repapa paradoxa* | 11 | 22 | (Gorochov and Warchałowska-Śliwa 2004) |
| Gryllidae | *Vasilia vietnamensis* | 17 | 20 | (Gorochov and Warchałowska-Śliwa 2004) |
| Gryllidae | *Neometrypus badius* | 14 | 28 | (Mesa and Garcia-Novo 2001) |
| Gryllidae | *Euscyrtus hemelytrus* | 17 | 17 | (Hewitt 1979) |
| Gryllidae | *Euscyrtus hemelytrus* | 23 | 23 | (Hewitt 1979) |
| Gryllidae | Oecanthus valensis | 18 | 22 | (Milach et al. 2016) |
| Phalangopsidae | *Strinatia brevinpennis* | 11 | 11 | (Mesa et al. 1999) |
| Phalangopsidae | *Luzaridella susurra* | 12 | 16 | (Timm et al. 2021) |

| | | | | |
|---|---|---|---|---|
| Phalangopsidae | *Luzarida lata* | 17 | 27 | (Timm et al. 2021) |
| Phalangopsidae | *Melanotes ornata* | 13 | 24 | (Timm et al. 2021) |
| Phalangopsidae | *Izecksohniella puri* | 11 | 20 | (Timm et al. 2021) |
| Phalangopsidae | *Aracamby mucuriensis* | 13 | 22 | (Timm et al. 2021) |
| Phalangopsidae | *Aracamby balneatorius* | 15 | 22 | (Timm et al. 2021) |
| Phalangopsidae | *Aracamby picinguabensis* | 17 | 20 | (Timm et al. 2021) |
| Phalangopsidae | *Strinatia brevipennis* | 13 | 24 | (Timm et al. 2021) |
| Phalangopsidae | *Strinatia teresopolis* | 11 | 20 | (Timm et al. 2021) |
| Phalangopsidae | *Ubiquepuella telytokous* | 17 | 33 | (Timm et al. 2021) |
| Phalangopsidae | *Ubiquepuella telytokous* | 17 | 35 | (Timm et al. 2021) |
| Phalangopsidae | *Adelosgryllus cruscastaneus* | 17 | 20 | (Timm et al. 2021) |
| Phalangopsidae | *Adelosgryllus similis* | 17 | 20 | (Timm et al. 2021) |
| Phalangopsidae | *Adelosgryllus rubricephalus* | 17 | 20 | (Timm et al. 2021) |
| Phalangopsidae | *Adelosgryllus rubricephalus* | 19 | 20 | (Timm et al. 2021) |
| Phalangopsidae | *Laranda meridionalis* | 21 | 36 | (Timm et al. 2021) |
| Phalangopsidae | *Endecous chape* | 19 | 32 | (Timm et al. 2021) |
| Phalangopsidae | *Endecous onthophagus* | 19 | 34 | (Timm et al. 2021) |
| Phalangopsidae | *Endecous onthophagus* | 19 | 32 | (Timm et al. 2021) |
| Phalangopsidae | *Endecous itatibensis* | 19 | 34 | (Timm et al. 2021) |
| Phalangopsidae | *Endecous cavernicolus* | 21 | 26 | (Timm et al. 2021) |
| Phalangopsidae | *Endecous betariensis* | 21 | 30 | (Timm et al. 2021) |
| Phalangopsidae | *Endecous alejomesai* | 21 | 28 | (Timm et al. 2021) |
| Phalangopsidae | *Endecous didymus* | 21 | 38 | (Timm et al. 2021) |
| Phalangopsidae | *Endecous troglobius* | 21 | 38 | (Timm et al. 2021) |
| Phalangopsidae | *Endecous ubajarensis* | 14 | 24 | (Timm et al. 2021) |
| Phalangopsidae | *Eidmanacris meridionalis* | 11 | 20 | (Timm et al. 2021) |
| Phalangopsidae | *Eidmanacris septentrionalis* | 11 | 20 | (Timm et al. 2021) |
| Phalangopsidae | *Eidmanacris alboannulata* | 13 | 19 | (Timm et al. 2021) |
| Phalangopsidae | *Eidmanacris bidentata* | 11 | 20 | (Timm et al. 2021) |

| | | | | |
|---|---|---|---|---|
| Phalangopsidae | *Eidmanacris corumbatai* | 13 | 16 | (Timm et al. 2021) |
| Phalangopsidae | *Seychellesia sp.* | 21 | 21 | (Timm et al. 2021) |
| Phalangopsidae | *Meloimorpha japonica* | 21 | 21 | (Timm et al. 2021) |
| Schizodactylidae | *Schizodactylus montrosus* | 14 | 14 | (Mesa 1965) |
| Tettigoniidae | *Hemisaga albilinea* | 31 | 31 | (Warchałowska-Śliwa and Bugrov 1998) |
| Tettigoniidae | *Hemisaga allira* | 31 | 31 | (Warchałowska-Śliwa 1998) |
| Tettigoniidae | *Hemisaga baileyi* | 31 | 31 | (Warchałowska-Śliwa 1998) |
| Tettigoniidae | *Hemisaga lenticulata* | 31 | 31 | (Warchałowska-Śliwa 1998) |
| Tettigoniidae | *Hemisga lanceolata* | 31 | 31 | (Warchałowska-Śliwa 1998) |
| Tettigoniidae | *Hemisaga lunodota* | 31 | 31 | (Warchałowska-Śliwa 1998) |
| Tettigoniidae | *Hemisaga mullaya* | 31 | 31 | (Warchałowska-Śliwa 1998) |
| Tettigoniidae | *Hemisaga pericalles* | 31 | 31 | (Warchałowska-Śliwa 1998) |
| Tettigoniidae | *Hemisaga saussurei* | 31 | 31 | (Warchałowska-Śliwa 1998) |
| Tettigoniidae | *Hemisaga undulata* | 31 | 31 | (Warchałowska-Śliwa 1998) |
| Tettigoniidae | *Hemisaga venator* | 31 | 31 | (Warchałowska-Śliwa 1998) |
| Tettigoniidae | *Sciarasaga quadrata* | 31 | 31 | (Warchałowska-Śliwa 1998) |
| Tettigoniidae | *Pachysaga australis* | 31 | 31 | (Warchałowska-Śliwa 1998) |
| Tettigoniidae | *Pachysaga croceopteryx* | 31 | 31 | (Warchałowska-Śliwa 1998) |
| Tettigoniidae | *Pachysaga eneabba* | 31 | 31 | (Warchałowska-Śliwa 1998) |
| Tettigoniidae | *Pachysaga munggai* | 31 | 31 | (Warchałowska-Śliwa 1998) |
| Tettigoniidae | *Pachysaga ocrocercus* | 31 | 31 | (Warchałowska-Śliwa 1998) |
| Tettigoniidae | *Psacadonotus insulanus* | 31 | 31 | (Warchałowska-Śliwa 1998) |
| Tettigoniidae | *Psacadonotus robustus* | 31 | 31 | (Warchałowska-Śliwa 1998) |
| Tettigoniidae | *Psacadonotus kenkulun* | 29 | 30 | (Warchałowska-Śliwa 1998) |
| Tettigoniidae | *Psacadonotus psithryros* | 29 | 30 | (Warchałowska-Śliwa 1998) |
| Tettigoniidae | *Psacadonotus viridis* | 29 | 30 | (Warchałowska-Śliwa 1998) |
| Tettigoniidae | *Baetica ustulata* | 25 | 29 | (Warchałowska-Śliwa 1998) |
| Tettigoniidae | *Bradyporus dasypus* | 27 | 32 | (Warchałowska-Śliwa 1998) |
| Tettigoniidae | *Bradyporus macrogaster pancici* | 25 | 25 | (Warchałowska-Śliwa 1998) |

| | | | | |
|---|---|---|---|---|
| Tettigoniidae | *Bradyporus macrogaster pancici* | 29 | 29 | (Warchałowska-Śliwa 1998) |
| Tettigoniidae | *Bradyporus macrogaster macrogaster* | 27 | 32 | (Warchałowska-Śliwa et al. 2013b) |
| Tettigoniidae | *Bradyporus macrogaster longicollis* | 27 | 32 | (Warchałowska-Śliwa et al. 2013) |
| Tettigoniidae | *Bradyporus onuscus* | 27 | 32 | (Warchałowska-Śliwa et al. 2013) |
| Tettigoniidae | *Callicrana seoanei* | 22 | 24 | (Warchałowska-Śliwa 1998) |
| Tettigoniidae | *Deracantha onos* | 29 | 29 | (Warchałowska-Śliwa 1998) |
| Tettigoniidae | *Deracanthina deracanthoides* | 31 | 31 | (Warchałowska-Śliwa 1998) |
| Tettigoniidae | *Deracanthella verrucosa* | 31 | 31 | (Warchałowska-Śliwa 1998) |
| Tettigoniidae | *Ephippiger cavannai* | 29 | 31 | (Warchałowska-Śliwa 1998) |
| Tettigoniidae | *Ephippiger discoidalis* | 29 | 31 | (Warchałowska-Śliwa 1998) |
| Tettigoniidae | *Ephippiger ephippiger vitium* | 29 | 31 | (Warchałowska-Śliwa 1998) |
| Tettigoniidae | *Ephippiger ephippiger* | 29 | 31 | (Warchałowska-Śliwa 1998) |
| Tettigoniidae | *Ephippiger ruffoi* | 29 | 31 | (Warchałowska-Śliwa 1998) |
| Tettigoniidae | *Ephippiger zelleri* | 29 | 31 | (Warchałowska-Śliwa 1998) |
| Tettigoniidae | *Ephippigerida nogrosignata* | 29 | 31 | (Warchałowska-Śliwa 1998) |
| Tettigoniidae | *Pycnogaster finotti* | 29 | 31 | (Warchałowska-Śliwa 1998) |
| Tettigoniidae | *Pycnogaster graellsii* | 29 | 31 | (Warchałowska-Śliwa 1998) |
| Tettigoniidae | *Pycnogaster inermis* | 29 | 31 | (Warchałowska-Śliwa 1998) |
| Tettigoniidae | *Pycnogaster sanchezgamezi* | 29 | 31 | (Warchałowska-Śliwa 1998) |
| Tettigoniidae | *Caraecercus fuscipennis* | 27 | 29 | (Warchałowska-Śliwa 1998) |
| Tettigoniidae | *Pycnogaster cucullata* | 28 | 31 | (Warchałowska-Śliwa 1998) |
| Tettigoniidae | *Pycnogaster cucullata* | 29 | 31 | (Warchałowska-Śliwa 1998) |
| Tettigoniidae | *Steropleurus stalii* | 29 | 31 | (Warchałowska-Śliwa et al. 2013) |
| Tettigoniidae | *Steropleurus pseudolus* | 27 | 31 | (Warchałowska-Śliwa et al. 2013) |
| Tettigoniidae | *Uromenus cockerelli* | 29 | 31 | (Warchałowska-Śliwa 1998) |
| Tettigoniidae | *Uromenus martorelli* | 27 | 29 | (Warchałowska-Śliwa 1998) |
| Tettigoniidae | *Uromenus martorelli* | 25 | 27 | (Warchałowska-Śliwa 1998) |
| Tettigoniidae | *Uromenus martorelli* | 23 | 27 | (Warchałowska-Śliwa 1998) |
| Tettigoniidae | *Uromenus brevicolis insularis* | 23 | 27 | (Warchałowska-Śliwa 1998) |

| | | | | |
|---|---|---|---|---|
| Tettigoniidae | *Uromenus elegans* | 23 | 27 | (Warchałowska-Śliwa 1998) |
| Tettigoniidae | *Uromenus riggioi* | 23 | 27 | (Warchałowska-Śliwa 1998) |
| Tettigoniidae | *Zychia varanovi* | 31 | 31 | (Warchałowska-Śliwa 1998) |
| Tettigoniidae | *Anelytra sp.* | 29 | 32 | (Warchałowska-Śliwa 1998) |
| Tettigoniidae | *Anelytra (Perianelytra) propria* | 27 | 30 | (Warchałowska-Sliwa and Bugrov 2000) |
| Tettigoniidae | *Conocephalus longipennis* | 33 | 33 | (Warchałowska-Śliwa 1998) |
| Tettigoniidae | *Conocephalus japonicus* | 33 | 33 | (Warchałowska-Śliwa 1998) |
| Tettigoniidae | *Conocephalus chinensis* | 33 | 33 | (Warchałowska-Śliwa 1998) |
| Tettigoniidae | *Conocephalus discolor* | 33 | 42 | (Warchałowska-Śliwa 1998) |
| Tettigoniidae | *Conocephalus dorsalis* | 33 | 42 | (Warchałowska-Śliwa 1998) |
| Tettigoniidae | *Conocephalus fasciatus* | 33 | 33 | (Warchałowska-Śliwa 1998) |
| Tettigoniidae | *Conocephalus gladiatus* | 31 | 31 | (Warchałowska-Śliwa 1998) |
| Tettigoniidae | *Conocephalus gladiatus* | 33 | 33 | (Warchałowska-Śliwa 1998) |
| Tettigoniidae | *Conocephalus melanum* | 33 | 33 | (Warchałowska-Śliwa 1998) |
| Tettigoniidae | *Conocephalus sp.* | 33 | 38 | (Warchałowska-Śliwa 1998) |
| Tettigoniidae | *Chortoscirtes serengeti* | 33 | 34 | (Hemp et al. 2013a) |
| Tettigoniidae | *Euconocephalus incertus* | 21 | 30 | (Warchałowska-Śliwa 1998) |
| Tettigoniidae | *Euconocephalus incertus* | 25 | 30 | (Warchałowska-Śliwa 1998) |
| Tettigoniidae | *Euconocephalus nasutus* | 21 | 30 | (Warchałowska-Śliwa 1998) |
| Tettigoniidae | *Euconocephalus pallidus* | 21 | 28 | (Warchałowska-Śliwa 1998) |
| Tettigoniidae | *Euconocephalus varius* | 21 | 30 | (Warchałowska-Śliwa 1998) |
| Tettigoniidae | *Hexacentrus inflatissimus* | 33 | 34 | (Gorochov and Warchałowska-Sliwa 2000) |
| Tettigoniidae | *Hexacentrus japonica hareyamai* | 33 | 33 | (Warchałowska-Śliwa 1998) |
| Tettigoniidae | *Hexacentrus unicolor* | 32 | 34 | (Warchałowska-Śliwa 1998) |
| Tettigoniidae | *Hexacentrus japonicus* | 31 | 31 | (Warchałowska-Śliwa 1998) |
| Tettigoniidae | *Hexacentrus mundus* | 31 | 32 | (Warchałowska-Śliwa 1998) |
| Tettigoniidae | *Genus A sp.* | 29 | 32 | (Warchałowska-Śliwa 1998) |
| Tettigoniidae | *Neoconocephalus infuscatus* | 27 | 27 | (Warchałowska-Śliwa 1998) |
| Tettigoniidae | *Neoconocephalus sp.* | 23 | 32 | (Warchałowska-Śliwa 1998) |

| | | | | |
|---|---|---|---|---|
| Tettigoniidae | *Liara tramlapensis* | 29 | 32 | (Gorochov and Warchałowska-Sliwa 2000) |
| Tettigoniidae | *Liaromorpha buonluoiensis* | 33 | 34 | (Gorochov and Warchałowska-Sliwa 2000) |
| Tettigoniidae | *Orchelimun concinnum* | 33 | 33 | (Warchałowska-Śliwa 1998) |
| Tettigoniidae | *Orchelimun vulgare* | 33 | 33 | (Warchałowska-Śliwa 1998) |
| Tettigoniidae | *Oxyprora flavicornis* | 29 | 29 | (Warchałowska-Śliwa 1998) |
| Tettigoniidae | *Phlesirtes kibonotensis* | 33 | 34 | (Hemp et al. 2010) |
| Tettigoniidae | *Phlesirtes kilimandjaricus* | 33 | 34 | (Hemp et al. 2010) |
| Tettigoniidae | *Phlesirtes meromontanus* | 33 | 34 | (Hemp et al. 2010) |
| Tettigoniidae | *Pseudorhynchus japonicus* | 29 | 32 | (Warchałowska-Śliwa 1998) |
| Tettigoniidae | *Ruspolia lineosus* | 25 | 25 | (Warchałowska-Śliwa 1998) |
| Tettigoniidae | *Ruspolia nitidula* | 21 | 21 | (Warchałowska-Śliwa 1998) |
| Tettigoniidae | *Sialaiana transiens* | 29 | 34 | (Gorochov and Warchałowska-Sliwa 2000) |
| Tettigoniidae | *Xesthophrys harvathi* | 29 | 32 | (Warchałowska-Śliwa 1998) |
| Tettigoniidae | *Acanthoplus speiseri* | 25 | 25 | (Warchałowska-Śliwa and Bugrov 2009) |
| Tettigoniidae | *Eugaster fernandezi* | 29 | 29 | (Warchałowska-Śliwa 1998) |
| Tettigoniidae | *Eugaster guyoni* | 29 | 29 | (Warchałowska-Śliwa 1998) |
| Tettigoniidae | *Eugaster spinulosa* | 29 | 29 | (Warchałowska-Śliwa 1998) |
| Tettigoniidae | *Hetrodes pupus* | 27 | 27 | (Warchałowska-Śliwa and Bugrov 2009) |
| Tettigoniidae | *Spalacomimus talpa* | 24 | 26 | (Warchałowska-Śliwa et al. 2015) |
| Tettigoniidae | *Spalacomimus verruciferus* | 24 | 26 | (Warchałowska-Śliwa et al. 2015) |
| Tettigoniidae | *Aerotegmina kilimandjarica* | 33 | 33 | (Hemp et al. 2013b) |
| Tettigoniidae | *Aerotegmin shengenae* | 27 | 34 | (Hemp et al. 2013b) |
| Tettigoniidae | *Euhexacentrus annulicornis* | 12 | 24 | (Warchałowska-Śliwa 1998) |
| Tettigoniidae | *Chlorobalius leucoviridis* | 20 | 21 | (Warchałowska-Śliwa 1998) |
| Tettigoniidae | *Yorkiella sp. 1* | 31 | 32 | (Warchałowska-Śliwa 1998) |
| Tettigoniidae | *Yorkiella sp. 2* | 31 | 32 | (Warchałowska-Śliwa 1998) |
| Tettigoniidae | *Kuzicus suzukii* | 31 | 31 | (Warchałowska-Śliwa 1998) |
| Tettigoniidae | *Leptoteratura albicorne* | 31 | 31 | (Warchałowska-Śliwa 1998) |
| Tettigoniidae | *Meconema thalassinum* | 27 | 29 | (Warchałowska-Śliwa 1998) |

| | | | | |
|---|---|---|---|---|
| Tettigoniidae | *Phisis sp.* | 21 | 25 | (Warchałowska-Śliwa 1998) |
| Tettigoniidae | *Xiphidiopsis straminula* | 33 | 36 | (Warchałowska-Śliwa 1998) |
| Tettigoniidae | *Xiphidiopsis sp.* | 33 | 46 | (Warchałowska-Śliwa 1998) |
| Tettigoniidae | *Xiphidiopsis lita* | 26 | 30 | (Warchałowska-Śliwa 1998) |
| Tettigoniidae | *Mecopoda elongata* | 29 | 57 | (Bugrov et al. 2004) |
| Tettigoniidae | *Mecopoda sp.* | 27 | 54 | (Warchałowska-Śliwa 1998) |
| Tettigoniidae | *Philoscirtus viridulus* | 29 | 38 | (Warchałowska-Śliwa et al. 2015) |
| Tettigoniidae | *Saga campbelli campbelli* | 27 | 32 | (Warchałowska-Śliwa et al. 2009) |
| Tettigoniidae | *Saga campbelli gracilis* | 27 | 32 | (Warchałowska-Śliwa et al. 2009) |
| Tettigoniidae | *Saga cappadocia* | 31 | 31 | (Warchałowska-Śliwa 1998) |
| Tettigoniidae | *Saga ephippigera* | 31 | 32 | (Warchałowska-Śliwa 1998) |
| Tettigoniidae | *Saga ephippigera* | 33 | 34 | (Warchałowska-Śliwa 1998) |
| Tettigoniidae | *Saga hellenica* | 29 | 32 | (Warchałowska-Śliwa 1998) |
| Tettigoniidae | *Saga natoliae* | 29 | 32 | (Warchałowska-Śliwa et al. 2009) |
| Tettigoniidae | *Saga ornata* | 31 | 32 | (Warchałowska-Śliwa 1998) |
| Tettigoniidae | *Saga rammei* | 23 | 28 | (Warchałowska-Śliwa et al. 2009) |
| Tettigoniidae | *Saga rhodiensis* | 29 | 32 | (Warchałowska-Śliwa et al. 2009) |
| Tettigoniidae | *Acrometopa servillea* | 31 | 31 | (Grzywacz et al. 2014b) |
| Tettigoniidae | *Amblycorypha rotundifolia* | 33 | 34 | (Warchałowska-Śliwa 1998) |
| Tettigoniidae | *Amblycorypha oblongifolia* | 31 | 32 | (Warchałowska-Śliwa 1998) |
| Tettigoniidae | *Amblycorypha oblongifolia* | 33 | 34 | (Warchałowska-Śliwa 1998) |
| Tettigoniidae | *Anaulacomera dimidiata* | 31 | 32 | (Warchałowska-Śliwa 1998) |
| Tettigoniidae | *Anaulacomera horti* | 31 | 32 | (Warchałowska-Śliwa 1998) |
| Tettigoniidae | *Anaulacomera sp.1* | 31 | 32 | (Warchałowska-Śliwa 1998) |
| Tettigoniidae | *Anaulacomera sp.2* | 31 | 32 | (Warchałowska-Śliwa 1998) |
| Tettigoniidae | *Anaulacomera sp.* | 31 | 32 | (Warchałowska-Śliwa 1998) |
| Tettigoniidae | *Ancistrura nigrovittata* | 31 | 31 | (Warchałowska-Śliwa 1998) |
| Tettigoniidae | *Andreiniimon nuptialis* | 31 | 31 | (Warchałowska-Śliwa 1998) |
| Tettigoniidae | *Anisotima japonica* | 27 | 28 | (Warchałowska-Śliwa 1998) |

| | | | | |
|---|---|---|---|---|
| Tettigoniidae | *Barbitistes constrictus* | 31 | 31 | (Warchałowska-Śliwa 1998) |
| Tettigoniidae | *Barbitistes serricauda* | 32 | 32 | (Warchałowska-Śliwa 1998) |
| Tettigoniidae | *Barbitistes kaltenbachi* | 31 | 31 | (Warchałowska-Śliwa 1998) |
| Tettigoniidae | *Barbitistes ocskayi* | 31 | 31 | (Warchałowska-Śliwa et al. 2013a) |
| Tettigoniidae | *Barbitistes yersini* | 31 | 31 | (Warchałowska-Śliwa et al. 2013b) |
| Tettigoniidae | *Caedicia marginata* | 20 | 21 | (Warchałowska-Śliwa 1998) |
| Tettigoniidae | *Caedicia sp. a* | 21 | 21 | (Warchałowska-Śliwa 1998) |
| Tettigoniidae | *Caedicia sp. b* | 21 | 21 | (Warchałowska-Śliwa 1998) |
| Tettigoniidae | *Caedicia sp. c* | 21 | 21 | (Warchałowska-Śliwa 1998) |
| Tettigoniidae | *Chloroscirtus forcipatus* | 31 | 31 | (Warchałowska-Śliwa 1998) |
| Tettigoniidae | *Cnemidophyllum citrifolium* | 25 | 25 | (Warchałowska-Śliwa 1998) |
| Tettigoniidae | *Dichopetala brevihastata* | 23 | 23 | (Warchałowska-Śliwa 1998) |
| Tettigoniidae | *Dichopetala tauriformis* | 23 | 23 | (Warchałowska-Śliwa 1998) |
| Tettigoniidae | *Dichopetala tauriformis* | 27 | 28 | (Warchałowska-Śliwa 1998) |
| Tettigoniidae | *Dichopetala femorata* | 17 | 17 | (Warchałowska-Śliwa 1998) |
| Tettigoniidae | *Dichopetala mexicana* | 17 | 17 | (Warchałowska-Śliwa 1998) |
| Tettigoniidae | *Diplophyllus acreanus* | 31 | 31 | (Warchałowska-Śliwa 1998) |
| Tettigoniidae | *Ducetia japonica* | 27 | 27 | (Warchałowska-Śliwa 1998) |
| Tettigoniidae | *Ducetia japonica* | 29 | 29 | (Warchałowska-Śliwa 1998) |
| Tettigoniidae | *Dysonia elegans* | 31 | 31 | (Warchałowska-Śliwa 1998) |
| Tettigoniidae | *Elimaea fallax* | 29 | 29 | (Warchałowska-Śliwa 1998) |
| Tettigoniidae | *Elimaea sp.* | 29 | 29 | (Warchałowska-Śliwa 1998) |
| Tettigoniidae | *Elimaea sucurigera* | 25 | 28 | (Warchałowska-Śliwa 1998) |
| Tettigoniidae | *Elimaea sucurigera* | 27 | 27 | (Warchałowska-Śliwa 1998) |
| Tettigoniidae | *Elimaea sucurigera* | 27 | 28 | (Warchałowska-Śliwa 1998) |
| Tettigoniidae | *Euceraia insignis* | 31 | 31 | (Warchałowska-Śliwa 1998) |
| Tettigoniidae | *Eurycorypha combretoides n. sp.* | 29 | 29 | (Hemp et al. 2013a) |
| Tettigoniidae | *E. meruensis* | 29 | 29 | (Hemp et al. 2013a) |
| Tettigoniidae | *E. punctipennis* | 29 | 30 | (Hemp et al. 2013a) |

| | | | | |
|---|---|---|---|---|
| Tettigoniidae | *E. resonans n. sp.* | 29 | 29 | (Hemp et al. 2013a) |
| Tettigoniidae | *E. varia* | 29 | 29 | (Hemp et al. 2013a) |
| Tettigoniidae | *Himertula kinneari* | 31 | 31 | (Warchałowska-Śliwa 1998) |
| Tettigoniidae | *Himertula sp.* | 31 | 31 | (Warchałowska-Śliwa 1998) |
| Tettigoniidae | *Holochlora japonica* | 31 | 31 | (Warchałowska-Śliwa 1998) |
| Tettigoniidae | *Holochlora spectabilis* | 29 | 29 | (Warchałowska-Śliwa 1998) |
| Tettigoniidae | *Holochlora spectabilis* | 31 | 31 | (Warchałowska-Śliwa 1998) |
| Tettigoniidae | *Holochlora sp.* | 31 | 31 | (Warchałowska-Śliwa 1998) |
| Tettigoniidae | *Hyperophora angustipennis* | 31 | 31 | (Warchałowska-Śliwa 1998) |
| Tettigoniidae | *Insara gracillima* | 31 | 31 | (Warchałowska-Śliwa 1998) |
| Tettigoniidae | *Insara tolteca* | 31 | 31 | (Warchałowska-Śliwa 1998) |
| Tettigoniidae | *Ischyra punctinervis* | 31 | 31 | (Hemp et al. 2013a) |
| Tettigoniidae | *Isophya hospodar* | 31 | 31 | (Grzywacz et al. 2014a) |
| Tettigoniidae | *Isophya straubei ssp.* | 31 | 31 | (Grzywacz et al. 2014a) |
| Tettigoniidae | *Isophya nervosa* | 31 | 31 | (Grzywacz et al. 2014a) |
| Tettigoniidae | *Isophya thracica* | 31 | 31 | (Grzywacz et al. 2014a) |
| Tettigoniidae | *Isophya stenocauda stenocauda* | 31 | 31 | (Grzywacz et al. 2014a) |
| Tettigoniidae | *Isophya stenocauda obenbergeri* | 31 | 31 | (Grzywacz et al. 2014a) |
| Tettigoniidae | *Isophya rectipennis* | 31 | 31 | (Grzywacz et al. 2014a) |
| Tettigoniidae | *Isophya pavelii* | 31 | 31 | (Grzywacz et al. 2014a) |
| Tettigoniidae | *Isophya bureschi* | 31 | 31 | (Grzywacz et al. 2014a) |
| Tettigoniidae | *Isophya andreevae* | 31 | 31 | (Grzywacz et al. 2014a) |
| Tettigoniidae | *Isophya miksici* | 31 | 31 | (Grzywacz et al. 2014a) |
| Tettigoniidae | *Isophya plevnensis* | 31 | 31 | (Grzywacz et al. 2014a) |
| Tettigoniidae | *Isophya longicaudata adamovici* | 31 | 31 | (Grzywacz et al. 2014a) |
| Tettigoniidae | *Isophya longicaudata longicaudata* | 31 | 31 | (Grzywacz et al. 2014a) |
| Tettigoniidae | *Isophya rh. leonorae* | 31 | 31 | (Grzywacz et al. 2014a) |
| Tettigoniidae | *Isophya rh. petkovi* | 31 | 31 | (Grzywacz et al. 2014a) |
| Tettigoniidae | *Isophya rh. rhodopensis* | 31 | 31 | (Grzywacz et al. 2014a) |

| | | | | |
|---|---|---|---|---|
| Tettigoniidae | *Isophya yaraligozi* | 31 | 31 | (Grzywacz et al. 2014a) |
| Tettigoniidae | *Isophya tosevski* | 31 | 31 | (Grzywacz et al. 2014a) |
| Tettigoniidae | *Isophya taurica* | 31 | 31 | (Grzywacz et al. 2014a) |
| Tettigoniidae | *Isophya obtusa* | 31 | 31 | (Grzywacz et al. 2014a) |
| Tettigoniidae | *Isophya gulae* | 31 | 31 | (Grzywacz et al. 2014a) |
| Tettigoniidae | *Isophya obtuse* | 31 | 31 | (Grzywacz et al. 2014a) |
| Tettigoniidae | *Isophya comptoxypha* | 31 | 31 | (Grzywacz et al. 2014a) |
| Tettigoniidae | *Isophya altaica* | 31 | 31 | (Grzywacz et al. 2014a) |
| Tettigoniidae | *Isophya brunneri* | 31 | 31 | (Grzywacz et al. 2014a) |
| Tettigoniidae | *Isophya modestior* | 31 | 31 | (Grzywacz et al. 2014a) |
| Tettigoniidae | *Isophya kraussii* | 31 | 31 | (Grzywacz et al. 2014a) |
| Tettigoniidae | *Isophya pienensis* | 31 | 31 | (Grzywacz et al. 2014a) |
| Tettigoniidae | *Isophya sp.* | 31 | 31 | (Grzywacz et al. 2014a) |
| Tettigoniidae | *Isophya zernovi* | 31 | 31 | (Grzywacz et al. 2014a) |
| Tettigoniidae | *Isophya autumnalis* | 31 | 31 | (Grzywacz et al. 2014a) |
| Tettigoniidae | *Isophya cf. amena* | 31 | 31 | (Grzywacz et al. 2014a) |
| Tettigoniidae | *Isophya schneideri* | 31 | 31 | (Grzywacz et al. 2014a) |
| Tettigoniidae | *Isophya sureyai* | 31 | 31 | (Grzywacz et al. 2014a) |
| Tettigoniidae | *Isophya aff. sureyai* | 31 | 31 | (Grzywacz et al. 2014a) |
| Tettigoniidae | *Isophya speciosa* | 31 | 31 | (Grzywacz et al. 2014a) |
| Tettigoniidae | *Isophya amplipennis* | 31 | 31 | (Grzywacz et al. 2014a) |
| Tettigoniidae | *Isophya rizeensis* | 31 | 31 | (Grzywacz et al. 2014a) |
| Tettigoniidae | *Isophya major* | 31 | 31 | (Grzywacz et al. 2014a) |
| Tettigoniidae | *Isophya pravdini pravdini* | 31 | 31 | (Warchałowska-Śliwa et al. 2008) |
| Tettigoniidae | *Isophya pravdini adamovici* | 31 | 31 | (Warchałowska-Śliwa et al. 2008) |
| Tettigoniidae | *Isophya modesta longicauda* | 31 | 31 | (Warchałowska-Śliwa et al. 2008) |
| Tettigoniidae | *Isophya kisi* | 31 | 31 | (Warchałowska-Śliwa et al. 2008) |
| Tettigoniidae | *Isophya brevipennis* | 31 | 31 | (Warchałowska-Śliwa 1998) |
| Tettigoniidae | *Isophya hemiptera* | 30 | 31 | (Warchałowska-Śliwa 1998) |

| | | | | |
|---|---|---|---|---|
| Tettigoniidae | *Isophya kalishevskii* | 31 | 31 | (Warchałowska-Śliwa 1998) |
| Tettigoniidae | *Isopsera sp.* | 32 | 33 | (Warchałowska-Śliwa 1998) |
| Tettigoniidae | *Itarissa sp.* | 17 | 21 | (Warchałowska-Śliwa 1998) |
| Tettigoniidae | *Kuwayamaea sapporensis* | 27 | 27 | (Warchałowska-Śliwa 1998) |
| Tettigoniidae | *Letana nigtosparsa* | 29 | 29 | (Warchałowska-Śliwa 1998) |
| Tettigoniidae | *Letana atomifera* | 27 | 29 | (Warchałowska-Śliwa 1998) |
| Tettigoniidae | *Leptophyes albivittata* | 29 | 29 | (Warchałowska-Śliwa 1998) |
| Tettigoniidae | *Leptophyes boscii* | 29 | 29 | (Warchałowska-Śliwa 1998) |
| Tettigoniidae | *Leptophyes discoidalis* | 31 | 31 | (Warchałowska-Śliwa 1998) |
| Tettigoniidae | *Leptophyes punctactissima* | 31 | 31 | (Warchałowska-Śliwa and Heller 1998) |
| Tettigoniidae | *Leptophyes punctactissima* | 31 | 32 | (Warchałowska-Śliwa and Heller 1998) |
| Tettigoniidae | *Leptophyes punctactissima* | 32 | 32 | (Warchałowska-Śliwa and Heller 1998) |
| Tettigoniidae | *Leptophyes laticauda* | 31 | 31 | (Warchałowska-Śliwa and Heller 1998) |
| Tettigoniidae | *Lunidia viridis* | 31 | 31 | (Hemp et al. 2010b) |
| Tettigoniidae | *Mendesius albosignatus* | 28 | 29 | (Warchałowska-Śliwa 1998) |
| Tettigoniidae | *Metaplastes oertzeni* | 31 | 31 | (Warchałowska-Śliwa et al. 2013b) |
| Tettigoniidae | *Metaplastes ornatus* | 31 | 31 | (Warchałowska-Śliwa 1998) |
| Tettigoniidae | *Microcentrum bicentenearia* | 31 | 31 | (Warchałowska-Śliwa 1998) |
| Tettigoniidae | *Microcentrum lanceolatum* | 31 | 31 | (Warchałowska-Śliwa 1998) |
| Tettigoniidae | *Microcentrum myrtifolium* | 31 | 31 | (Warchałowska-Śliwa 1998) |
| Tettigoniidae | *Microcentrum sp.* | 31 | 31 | (Warchałowska-Śliwa 1998) |
| Tettigoniidae | *Microcentrum sp.* | 33 | 33 | (Warchałowska-Śliwa 1998) |
| Tettigoniidae | *Odontura aspericauda* | 31 | 31 | (Grzywacz et al. 2014a) |
| Tettigoniidae | *Odontura aspericauda* | 28 | 31 | (Grzywacz et al. 2014a) |
| Tettigoniidae | *Odontura aspericauda* | 31 | 32 | (Grzywacz et al. 2014a) |
| Tettigoniidae | *Odontura arcuata* | 28 | 28 | (Warchałowska-Śliwa 1998) |
| Tettigoniidae | *Odontura stenoxypha* | 28 | 29 | (Warchałowska-Śliwa 1998) |
| Tettigoniidae | *Odontura stenoxypha* | 28 | 28 | (Warchałowska-Śliwa 1998) |
| Tettigoniidae | *Odontura calaritana* | 27 | 27 | (Warchałowska-Śliwa 1998) |

| | | | | |
|---|---|---|---|---|
| Tettigoniidae | *Odontura glabircauda* | 27 | 28 | (Grzywacz et al. 2014a) |
| Tettigoniidae | *Odontura macphersoni* | 29 | 30 | (Grzywacz et al. 2014a) |
| Tettigoniidae | *Odontura maroccana* | 27 | 27 | (Warchałowska-Śliwa 1998) |
| Tettigoniidae | *Scaphura nigra* | 26 | 29 | (Mesa et al. 2010) |
| Tettigoniidae | *Stilpnochlora azteca* | 33 | 33 | (Warchałowska-Śliwa 1998) |
| Tettigoniidae | *Stilpnochlora marginella* | 33 | 33 | (Warchałowska-Śliwa 1998) |
| Tettigoniidae | *Stilpnochlora quadrata* | 31 | 31 | (Warchałowska-Śliwa 1998) |
| Tettigoniidae | *Parableta affinis* | 31 | 31 | (Warchałowska-Śliwa 1998) |
| Tettigoniidae | *Paratota ferrerai* | 31 | 31 | (Warchałowska-Śliwa 1998) |
| Tettigoniidae | *Pelerinus alienus* | 29 | 29 | (Warchałowska-Śliwa 1998) |
| Tettigoniidae | *Phaneroptera exigua* | 31 | 31 | (Warchałowska-Śliwa 1998) |
| Tettigoniidae | *Phaneroptera falcata* | 27 | 27 | (Warchałowska-Śliwa 1998) |
| Tettigoniidae | *Phaneroptera nigroantennata* | 25 | 25 | (Warchałowska-Śliwa 1998) |
| Tettigoniidae | *Phaneroptera nigroantennata* | 27 | 27 | (Warchałowska-Śliwa 1998) |
| Tettigoniidae | *Phaneroptera nana nana* | 27 | 27 | (Warchałowska-Śliwa 1998) |
| Tettigoniidae | *Phaneroptera quinquisignata* | 29 | 30 | (Warchałowska-Śliwa 1998) |
| Tettigoniidae | *Phaneroptera quinquisignata* | 29 | 31 | (Warchałowska-Śliwa 1998) |
| Tettigoniidae | *Phaneropterops piracicabensis* | 29 | 29 | (Warchałowska-Śliwa 1998) |
| Tettigoniidae | *Phonochorion artvinensis* | 31 | 31 | (Warchałowska-Śliwa et al. 2013b) |
| Tettigoniidae | *Philophyllia guttulata* | 31 | 31 | (Warchałowska-Śliwa et al. 2000) |
| Tettigoniidae | *Phrixa nasuta* | 25 | 25 | (Warchałowska-Śliwa et al. 2000) |
| Tettigoniidae | *Poecilimon lederei* | 31 | 31 | (Grzywacz et al. 2014a) |
| Tettigoniidae | *Poecilimon aff. lederei* | 31 | 31 | (Grzywacz et al. 2014a) |
| Tettigoniidae | *Poecilimon orbelicus* | 31 | 31 | (Grzywacz et al. 2014a) |
| Tettigoniidae | *Poecilimon armeniacus* | 31 | 31 | (Grzywacz et al. 2014a) |
| Tettigoniidae | *Poecilimon ampliatus* | 31 | 31 | (Grzywacz et al. 2014a) |
| Tettigoniidae | *Poecilimon pechevi* | 31 | 31 | (Grzywacz et al. 2014a) |
| Tettigoniidae | *Poecilimon ebneri* | 31 | 31 | (Grzywacz et al. 2014a) |
| Tettigoniidae | *Poecilimon klisiurensis* | 31 | 31 | (Grzywacz et al. 2014a) |

| | | | | |
|---|---|---|---|---|
| Tettigoniidae | *Poecilimon marmaraensis* | 31 | 31 | (Grzywacz et al. 2014a) |
| Tettigoniidae | *Poecilimon cf. karakushi* | 31 | 31 | (Grzywacz et al. 2014a) |
| Tettigoniidae | *Poecilimon erisi* | 31 | 31 | (Grzywacz et al. 2014a) |
| Tettigoniidae | *Poecilimon serratus* | 31 | 31 | (Grzywacz et al. 2014a) |
| Tettigoniidae | *Poecilimon toros* | 31 | 31 | (Grzywacz et al. 2014a) |
| Tettigoniidae | *Poecilimon cevus* | 31 | 31 | (Grzywacz et al. 2014a) |
| Tettigoniidae | *Poecilimon fussii* | 31 | 31 | (Grzywacz et al. 2014b) |
| Tettigoniidae | *Poecilimon pliginskii* | 31 | 31 | (Grzywacz et al. 2014a) |
| Tettigoniidae | *Poecilimon aff. bischoffi* | 31 | 31 | (Grzywacz et al. 2014a) |
| Tettigoniidae | *Poecilimon bischoffi* | 31 | 31 | (Grzywacz et al. 2014a) |
| Tettigoniidae | *Poecilimon bosphoricus* | 31 | 31 | (Grzywacz et al. 2014a) |
| Tettigoniidae | *Poecilimon miramae* | 31 | 31 | (Grzywacz et al. 2014a) |
| Tettigoniidae | *Poecilimon heinrichi* | 31 | 31 | (Grzywacz et al. 2014a) |
| Tettigoniidae | *Poecilimon roseoviridis* | 31 | 31 | (Grzywacz et al. 2014a) |
| Tettigoniidae | *Poecilimon turcicus* | 31 | 31 | (Grzywacz et al. 2014a) |
| Tettigoniidae | *Poecilimon anatolicus* | 31 | 31 | (Grzywacz et al. 2014a) |
| Tettigoniidae | *Poecilimon similis* | 31 | 31 | (Grzywacz et al. 2014a) |
| Tettigoniidae | *Poecilimon chopardi* | 31 | 31 | (Grzywacz et al. 2014a) |
| Tettigoniidae | *Poecilimon brunneri* | 31 | 31 | (Grzywacz et al. 2014a) |
| Tettigoniidae | *Poecilimon ukrainicus* | 31 | 31 | (Grzywacz et al. 2014a) |
| Tettigoniidae | *Poecilimon macedonicus* | 31 | 32 | (Grzywacz et al. 2014a) |
| Tettigoniidae | *Poecilimon zwicki* | 31 | 31 | (Grzywacz et al. 2014a) |
| Tettigoniidae | *Poecilimon jonicus* | 31 | 31 | (Grzywacz et al. 2014a) |
| Tettigoniidae | *Poecilimon martinae* | 31 | 32 | (Grzywacz et al. 2014a) |
| Tettigoniidae | *Poecilimon schimidtii* | 31 | 31 | (Grzywacz et al. 2014a) |
| Tettigoniidae | *Poecilimon zonatus* | 31 | 31 | (Grzywacz et al. 2014a) |
| Tettigoniidae | *Poecilimon jablanicensis* | 31 | 31 | (Grzywacz et al. 2014a) |
| Tettigoniidae | *Poecilimon ornatus* | 31 | 31 | (Grzywacz et al. 2014a) |
| Tettigoniidae | *Poecilimon affinis* | 31 | 31 | (Grzywacz et al. 2014a) |

| | | | | |
|---|---|---|---|---|
| Tettigoniidae | *Poecilimon ataturki* | 31 | 31 | (Grzywacz et al. 2014a) |
| Tettigoniidae | *Poecilimon celebi* | 31 | 31 | (Grzywacz et al. 2014a) |
| Tettigoniidae | *Poecilimon artedentatus* | 31 | 31 | (Warchałowska-Śliwa 1998) |
| Tettigoniidae | *Poecilimon heroicus* | 31 | 31 | (Warchałowska-Śliwa 1998) |
| Tettigoniidae | *Poecilimon hoelzeli* | 31 | 31 | (Warchałowska-Śliwa 1998) |
| Tettigoniidae | *Poecilimon intermedius* | 31 | 31 | (Warchałowska-Śliwa 1998) |
| Tettigoniidae | *Poecilimon jonicus tessellatus* | 31 | 32 | (Warchałowska-Śliwa 1998) |
| Tettigoniidae | *Poecilimon laevissimus* | 31 | 31 | (Warchałowska-Śliwa 1998) |
| Tettigoniidae | *Poecilimon mariannae* | 31 | 31 | (Warchałowska-Śliwa 1998) |
| Tettigoniidae | *Poecilimon nobilis* | 31 | 31 | (Warchałowska-Śliwa 1998) |
| Tettigoniidae | *Poecilimon zimmeri* | 31 | 31 | (Warchałowska-Śliwa 1998) |
| Tettigoniidae | *Poecilimon veluchianus veluchianus* | 31 | 31 | (Warchałowska-Śliwa 1998) |
| Tettigoniidae | *Poecilimon veluchianus minor* | 31 | 31 | (Warchałowska-Śliwa 1998) |
| Tettigoniidae | *Poecilimon unispinosus* | 31 | 31 | (Warchałowska-Śliwa 1998) |
| Tettigoniidae | *Poecilimon thoracicus* | 31 | 31 | (Warchałowska-Śliwa 1998) |
| Tettigoniidae | *Poecilimon thessalicus* | 31 | 31 | (Warchałowska-Śliwa 1998) |
| Tettigoniidae | *Poecilimon superbus* | 31 | 31 | (Warchałowska-Śliwa 1998) |
| Tettigoniidae | *Poecilimon scythicus* | 31 | 31 | (Warchałowska-Śliwa 1998) |
| Tettigoniidae | *Poecilimon propinquus* | 31 | 31 | (Warchałowska-Śliwa 1998) |
| Tettigoniidae | *Poecilimon aff. glandifer* | 31 | 31 | (Grzywacz et al. 2014a) |
| Tettigoniidae | *Poecilimon aegaeus* | 31 | 31 | (Warchałowska-Śliwa et al. 2000) |
| Tettigoniidae | *Poecilimon deplanatus* | 31 | 31 | (Warchałowska-Śliwa et al. 2000) |
| Tettigoniidae | *Poecilimon ikariensis* | 31 | 31 | (Warchałowska-Śliwa et al. 2000) |
| Tettigoniidae | *Polichne parvicauda* | 31 | 31 | (Warchałowska-Śliwa 1998) |
| Tettigoniidae | *Polysarcus denticauda* | 31 | 31 | (Warchałowska-Śliwa 1998) |
| Tettigoniidae | *Polysarcus zacharovi* | 31 | 31 | (Warchałowska-Śliwa 1998) |
| Tettigoniidae | *Polysarcus cf. elbursianus* | 31 | 31 | (Warchałowska-Śliwa et al. 2013a) |
| Tettigoniidae | *Polysarcus zigana* | 31 | 31 | (Warchałowska-Śliwa et al. 2013a) |
| Tettigoniidae | *Parapoecilimon antaltyaensis* | 31 | 31 | (Warchałowska-Śliwa et al. 2013a) |

| | | | | |
|---|---|---|---|---|
| Tettigoniidae | *Pycnopalpa bicordata* | 31 | 31 | (Warchałowska-Śliwa 1998) |
| Tettigoniidae | *Scudderia curvicanda* | 31 | 31 | (Warchałowska-Śliwa 1998) |
| Tettigoniidae | *Scudderia furcata furcata* | 31 | 31 | (Warchałowska-Śliwa 1998) |
| Tettigoniidae | *Scudderia texensis* | 31 | 31 | (Warchałowska-Śliwa 1998) |
| Tettigoniidae | *Scudderia psitillata* | 29 | 29 | (Warchałowska-Śliwa 1998) |
| Tettigoniidae | *Scudderia sp.* | 31 | 31 | (Warchałowska-Śliwa 1998) |
| Tettigoniidae | *Theudoria malanocnemis* | 30 | 31 | (Warchałowska-Śliwa 1998) |
| Tettigoniidae | *Tinzeda albosignata* | 25 | 25 | (Warchałowska-Śliwa 1998) |
| Tettigoniidae | *Topana aquilari* | 31 | 31 | (Warchałowska-Śliwa 1998) |
| Tettigoniidae | *Torbia viridissima* | 19 | 19 | (Warchałowska-Śliwa 1998) |
| Tettigoniidae | *Trigonocorypha unicolor* | 19 | 19 | (Warchałowska-Śliwa 1998) |
| Tettigoniidae | *Trigonocorypha unicolor* | 29 | 29 | (Warchałowska-Śliwa 1998) |
| Tettigoniidae | *Tylopsis lilifolia* | 31 | 31 | (Warchałowska-Śliwa 1998) |
| Tettigoniidae | *Viadana longicercata* | 29 | 29 | (Warchałowska-Śliwa 1998) |
| Tettigoniidae | *Bliastes viridifrons* | 33 | 36 | (Ferreira and Mesa 2010) |
| Tettigoniidae | *Diophanes amazonensis* | 29 | 34 | (Ferreira and Mesa 2010) |
| Tettigoniidae | *Diophanes scaberrimus* | 35 | 36 | (Ferreira and Mesa 2010) |
| Tettigoniidae | *Jamaicana flava* | 35 | 35 | (Ferreira and Mesa 2010) |
| Tettigoniidae | *Jamaicana unicolor* | 33 | 35 | (Ferreira and Mesa 2010) |
| Tettigoniidae | *Jamaicana subgutatta* | 33 | 35 | (Ferreira and Mesa 2010) |
| Tettigoniidae | *Leptotettix crassicerci* | 31 | 34 | (Ferreira and Mesa 2010) |
| Tettigoniidae | *Leptotettix humaita* | 35 | 36 | (Ferreira and Mesa 2010) |
| Tettigoniidae | *Meroncidius intermedius* | 31 | 35 | (Ferreira and Mesa 2010) |
| Tettigoniidae | *Phyllozelus pectinatus* | 35 | 36 | (Warchałowska-Śliwa 1998) |
| Tettigoniidae | *Sathrophylia sp.* | 35 | 35 | (Ferreira and Mesa 2010) |
| Tettigoniidae | *Sathrophylia rugosa* | 31 | 31 | (Ferreira and Mesa 2010) |
| Tettigoniidae | *Sathrophylia femorata* | 35 | 35 | (Ferreira and Mesa 2010) |
| Tettigoniidae | *Alticolana alticola* | 31 | 31 | (Warchałowska-Śliwa 1998) |
| Tettigoniidae | *Anabrus simplex* | 29 | 31 | (Warchałowska-Śliwa 1998) |

| | | | | |
|---|---|---|---|---|
| Tettigoniidae | *Anabrus sp.* | 33 | 33 | (Warchałowska-Śliwa 1998) |
| Tettigoniidae | *Anadrymadusa robusta* | 27 | 29 | (Warchałowska-Śliwa 1998) |
| Tettigoniidae | *Anadrymadusa picta* | 27 | 29 | (Warchałowska-Śliwa 1998) |
| Tettigoniidae | *Antipodectes brevicaudus* | 31 | 31 | (Warchałowska-Śliwa 1998) |
| Tettigoniidae | *Antipodectes giganteus* | 31 | 31 | (Warchałowska-Śliwa 1998) |
| Tettigoniidae | *Antipodectes graminicolus* | 29 | 29 | (Warchałowska-Śliwa 1998) |
| Tettigoniidae | *Anatlanticus koreanus* | 25 | 29 | (Warchałowska-Śliwa 1998) |
| Tettigoniidae | *Anterastes serbicus* | 31 | 31 | (Warchałowska-Śliwa et al. 2005) |
| Tettigoniidae | *Ateloplus hesperus* | 29 | 31 | (Warchałowska-Śliwa 1998) |
| Tettigoniidae | *Atlanticus brunneri* | 29 | 32 | (Warchałowska-Śliwa 1998) |
| Tettigoniidae | *Atlanticus pachymerus* | 25 | 30 | (Warchałowska-Śliwa 1998) |
| Tettigoniidae | *Atlanticus testaceus* | 25 | 30 | (Warchałowska-Śliwa 1998) |
| Tettigoniidae | *Bergiola montana* | 27 | 30 | (Warchałowska-Śliwa 1998) |
| Tettigoniidae | *Bucephaloptera bucephala* | 31 | 31 | (Warchałowska-Śliwa et al. 2005) |
| Tettigoniidae | *Capnobotes arizonensis* | 23 | 30 | (Warchałowska-Śliwa 1998) |
| Tettigoniidae | *Capnobotes attenuatus* | 23 | 30 | (Warchałowska-Śliwa 1998) |
| Tettigoniidae | *Capnobotes occidentalis* | 23 | 30 | (Warchałowska-Śliwa 1998) |
| Tettigoniidae | *Ceraeocercus fuscipennis* | 27 | 29 | (Warchałowska-Śliwa 1998) |
| Tettigoniidae | *Chinandectes neenan* | 29 | 29 | (Warchałowska-Śliwa 1998) |
| Tettigoniidae | *Chlorodectes laquax* | 27 | 30 | (Warchałowska-Śliwa 1998) |
| Tettigoniidae | *Chlorodectes baldersoni* | 25 | 28 | (Warchałowska-Śliwa 1998) |
| Tettigoniidae | *Chlorodectes montanus* | 25 | 28 | (Warchałowska-Śliwa 1998) |
| Tettigoniidae | *Chlorodectes ligaenus* | 23 | 26 | (Warchałowska-Śliwa 1998) |
| Tettigoniidae | *Clinopleura insfuscata* | 29 | 31 | (Warchałowska-Śliwa 1998) |
| Tettigoniidae | *Clinopleura minuta* | 29 | 31 | (Warchałowska-Śliwa 1998) |
| Tettigoniidae | *Ctenodecticus granatensis* | 26 | 33 | (Warchałowska-Śliwa 1998) |
| Tettigoniidae | *Ctenodecticus major* | 31 | 34 | (Warchałowska-Śliwa 1998) |
| Tettigoniidae | *Decticita brevicauda* | 31 | 31 | (Warchałowska-Śliwa 1998) |
| Tettigoniidae | *Decticus albifrons* | 31 | 31 | (Warchałowska-Śliwa 1998) |

| | | | | |
|---|---|---|---|---|
| Tettigoniidae | *Decticus verrucivorus* | 31 | 31 | (Warchałowska-Śliwa 1998) |
| Tettigoniidae | *Decticus verrucivorus* | 23 | 31 | (Warchałowska-Śliwa 1998) |
| Tettigoniidae | *Dexerra acanthiterga* | 29 | 29 | (Warchałowska-Śliwa 1998) |
| Tettigoniidae | *Dexerra serrata* | 29 | 29 | (Warchałowska-Śliwa 1998) |
| Tettigoniidae | *Dexerra turpis* | 29 | 29 | (Warchałowska-Śliwa 1998) |
| Tettigoniidae | *Dexerra vigescens* | 29 | 29 | (Warchałowska-Śliwa 1998) |
| Tettigoniidae | *Drymadusa dorsalis limbata* | 27 | 30 | (Warchałowska-Śliwa et al. 2005) |
| Tettigoniidae | *Drymadusella hissarica* | 31 | 31 | (Warchałowska-Śliwa 1998) |
| Tettigoniidae | *Ectopistidectes daptes* | 27 | 31 | (Warchałowska-Śliwa 1998) |
| Tettigoniidae | *Ectopistidectes viridis* | 27 | 31 | (Warchałowska-Śliwa 1998) |
| Tettigoniidae | *Eremopedes ephippita* | 31 | 31 | (Warchałowska-Śliwa 1998) |
| Tettigoniidae | *Eulithoxenus mongolicus* | 31 | 31 | (Warchałowska-Śliwa 1998) |
| Tettigoniidae | *Eupholidoptera anatolica* | 31 | 31 | (Warchałowska-Śliwa et al. 2005) |
| Tettigoniidae | *Eupholidoptera annulipes* | 31 | 31 | (Warchałowska-Śliwa et al. 2005) |
| Tettigoniidae | *Eupholidoptera epirotica* | 31 | 31 | (Warchałowska-Śliwa et al. 2005) |
| Tettigoniidae | *Eupholidoptera tauricola* | 31 | 31 | (Warchałowska-Śliwa et al. 2005) |
| Tettigoniidae | *Eupholidoptera smyrnensis* | 31 | 31 | (Warchałowska-Śliwa et al. 2005) |
| Tettigoniidae | *Eupholidoptera chabrieri garganica* | 31 | 31 | (Warchałowska-Śliwa et al. 2005) |
| Tettigoniidae | *Eupholidoptera megastyla* | 31 | 31 | (Warchałowska-Śliwa et al. 2005) |
| Tettigoniidae | *Eupholidoptera sp.* | 31 | 31 | (Warchałowska-Śliwa et al. 2005) |
| Tettigoniidae | *Eupholidoptera prasina* | 31 | 31 | (Warchałowska-Śliwa et al. 2005) |
| Tettigoniidae | *Eupholidoptera mersinensis* | 31 | 31 | (Warchałowska-Śliwa et al. 2005) |
| Tettigoniidae | *Eupholidoptera karabagi* | 31 | 31 | (Warchałowska-Śliwa et al. 2005) |
| Tettigoniidae | *Eupholidoptera icariensis* | 31 | 31 | (Warchałowska-Śliwa et al. 2005) |
| Tettigoniidae | *Gampsocleis burgeri* | 31 | 31 | (Warchałowska-Śliwa 1998) |
| Tettigoniidae | *Gampsocleis gratiosa* | 31 | 31 | (Warchałowska-Śliwa 1998) |
| Tettigoniidae | *Gampsocleis sedacovvii* | 31 | 31 | (Warchałowska-Śliwa 1998) |
| Tettigoniidae | *Gampsocleis sedacovvii sedacovii* | 31 | 31 | (Warchałowska-Śliwa 1998) |
| Tettigoniidae | *Gampsocleis ussuriensis* | 31 | 31 | (Warchałowska-Śliwa 1998) |

| | | | | |
|---|---|---|---|---|
| Tettigoniidae | *Gampsocleis ryukywensis* | 31 | 32 | (Warchałowska-Śliwa 1998) |
| Tettigoniidae | *Gampsocleis glabra* | 23 | 36 | (Warchałowska-Śliwa 1998) |
| Tettigoniidae | *Gampsocleis abbrevaita* | 23 | 36 | (Warchałowska-Śliwa 1998) |
| Tettigoniidae | *Glenbalodectes amaroo* | 33 | 33 | (Warchałowska-Śliwa 1998) |
| Tettigoniidae | *Glenbalodectes narraga* | 33 | 33 | (Warchałowska-Śliwa 1998) |
| Tettigoniidae | *Glenbalodectes norrisi* | 33 | 33 | (Warchałowska-Śliwa 1998) |
| Tettigoniidae | *Glyphonotus thoracicus* | 21 | 26 | (Warchałowska-Śliwa 1998) |
| Tettigoniidae | *Idiostatus aegualis* | 29 | 31 | (Warchałowska-Śliwa 1998) |
| Tettigoniidae | *Idiostatus apollo* | 29 | 31 | (Warchałowska-Śliwa 1998) |
| Tettigoniidae | *Idiostatus bechteli* | 29 | 31 | (Warchałowska-Śliwa 1998) |
| Tettigoniidae | *Idiostatus elegans* | 29 | 32 | (Warchałowska-Śliwa 1998) |
| Tettigoniidae | *Idiostatus fuscopunctatus* | 29 | 32 | (Warchałowska-Śliwa 1998) |
| Tettigoniidae | *Idiostatus gurneyi* | 29 | 31 | (Warchałowska-Śliwa 1998) |
| Tettigoniidae | *Idiostatus hermani* | 29 | 31 | (Warchałowska-Śliwa 1998) |
| Tettigoniidae | *Idiostatus inermis* | 29 | 31 | (Warchałowska-Śliwa 1998) |
| Tettigoniidae | *Idiostatus inermoides* | 29 | 32 | (Warchałowska-Śliwa 1998) |
| Tettigoniidae | *Idiostatus kothleenae* | 29 | 31 | (Warchałowska-Śliwa 1998) |
| Tettigoniidae | *Idiostatus magnificus* | 29 | 31 | (Warchałowska-Śliwa 1998) |
| Tettigoniidae | *Idiostatus rehni* | 29 | 32 | (Warchałowska-Śliwa 1998) |
| Tettigoniidae | *Idiostatus nevadensis* | 27 | 29 | (Warchałowska-Śliwa 1998) |
| Tettigoniidae | *Idionotus brunneus* | 27 | 29 | (Warchałowska-Śliwa 1998) |
| Tettigoniidae | *Idionotus tehachapi* | 27 | 29 | (Warchałowska-Śliwa 1998) |
| Tettigoniidae | *Ixalodectes nigrifrons* | 23 | 23 | (Warchałowska-Śliwa 1998) |
| Tettigoniidae | *Ixalodectes uptoni* | 23 | 23 | (Warchałowska-Śliwa 1998) |
| Tettigoniidae | *Ixalodectes megacercus* | 15 | 19 | (Warchałowska-Śliwa 1998) |
| Tettigoniidae | *Ixalodectes whitei* | 15 | 19 | (Warchałowska-Śliwa 1998) |
| Tettigoniidae | *Lanciana albidicornsi* | 25 | 25 | (Warchałowska-Śliwa 1998) |
| Tettigoniidae | *Lanciana montana* | 25 | 27 | (Warchałowska-Śliwa 1998) |
| Tettigoniidae | *Lanciana semilata* | 25 | 25 | (Warchałowska-Śliwa 1998) |

| | | | | |
|---|---|---|---|---|
| Tettigoniidae | *Lanciana hisperpotana* | 21 | 23 | (Warchałowska-Śliwa 1998) |
| Tettigoniidae | *Lanciana occidentalis* | 21 | 23 | (Warchałowska-Śliwa 1998) |
| Tettigoniidae | *Metaballus alatus* | 27 | 27 | (Warchałowska-Śliwa 1998) |
| Tettigoniidae | *Metaballus brevipennis* | 27 | 27 | (Warchałowska-Śliwa 1998) |
| Tettigoniidae | *Metaballus bynoei* | 27 | 27 | (Warchałowska-Śliwa 1998) |
| Tettigoniidae | *Metaballus dectiocoides* | 27 | 27 | (Warchałowska-Śliwa 1998) |
| Tettigoniidae | *Metaballus frontalis* | 27 | 27 | (Warchałowska-Śliwa 1998) |
| Tettigoniidae | *Metaballus litus* | 27 | 27 | (Warchałowska-Śliwa 1998) |
| Tettigoniidae | *Metaballus mesopterus* | 27 | 27 | (Warchałowska-Śliwa 1998) |
| Tettigoniidae | *Metaballus murunatus* | 27 | 27 | (Warchałowska-Śliwa 1998) |
| Tettigoniidae | *Metaballus nchigyues* | 27 | 27 | (Warchałowska-Śliwa 1998) |
| Tettigoniidae | *Metrioptera arnoldi* | 31 | 31 | (Warchałowska-Śliwa 1998) |
| Tettigoniidae | *Metrioptera bicolor* | 31 | 31 | Warchałowska-Śliwa et at. 2005 |
| Tettigoniidae | *Metrioptera bonetti* | 31 | 31 | (Warchałowska-Śliwa 1998) |
| Tettigoniidae | *Metrioptera brachyptera* | 31 | 31 | (Warchałowska-Śliwa 1998) |
| Tettigoniidae | *Metrioptera japonica* | 31 | 31 | (Warchałowska-Śliwa 1998) |
| Tettigoniidae | *Metrioptera oblongicollis* | 31 | 31 | (Warchałowska-Śliwa et al. 2005) |
| Tettigoniidae | *Metrioptera saussureana* | 29 | 31 | (Warchałowska-Śliwa 1998) |
| Tettigoniidae | *Metrioptera roeselii ambitiosa* | 31 | 31 | (Warchałowska-Śliwa et al. 2005) |
| Tettigoniidae | *Metrioptera roeselii roeselii* | 31 | 31 | (Warchałowska-Śliwa et al. 2005) |
| Tettigoniidae | *Metrioptera ussuriana* | 31 | 31 | (Warchałowska-Śliwa 1998) |
| Tettigoniidae | *Montana alexandra* | 31 | 32 | (Warchałowska-Śliwa 1998) |
| Tettigoniidae | *Montana everssanni* | 31 | 31 | (Warchałowska-Śliwa 1998) |
| Tettigoniidae | *Montana montana* | 31 | 31 | (Warchałowska-Śliwa 1998) |
| Tettigoniidae | *Montana daghestanica* | 29 | 31 | (Warchałowska-Śliwa 1998) |
| Tettigoniidae | *Montana tomini* | 29 | 34 | (Warchałowska-Śliwa 1998) |
| Tettigoniidae | *Nanodectes dooloides* | 23 | 23 | (Warchałowska-Śliwa 1998) |
| Tettigoniidae | *Nanodectes gladiator* | 21 | 21 | (Warchałowska-Śliwa 1998) |
| Tettigoniidae | *Nanodectes harpax* | 21 | 21 | (Warchałowska-Śliwa 1998) |

| | | | | |
|---|---|---|---|---|
| Tettigoniidae | *Nanodectes platycerus* | 21 | 21 | (Warchałowska-Śliwa 1998) |
| Tettigoniidae | *Nanodectes brachyrus* | 19 | 21 | (Warchałowska-Śliwa 1998) |
| Tettigoniidae | *Nanodectes bulbicerus* | 19 | 19 | (Warchałowska-Śliwa 1998) |
| Tettigoniidae | *Nanodectes dooloo* | 19 | 21 | (Warchałowska-Śliwa 1998) |
| Tettigoniidae | *Nanodectes veprephila* | 17 | 21 | (Warchałowska-Śliwa 1998) |
| Tettigoniidae | *Nanodectes triodiae* | 15 | 21 | (Warchałowska-Śliwa 1998) |
| Tettigoniidae | *Nanodectes triodiae* | 15 | 19 | (Warchałowska-Śliwa 1998) |
| Tettigoniidae | *Nanodectes triodiae* | 17 | 21 | (Warchałowska-Śliwa 1998) |
| Tettigoniidae | *Nanodectes triodiae* | 19 | 21 | (Warchałowska-Śliwa 1998) |
| Tettigoniidae | *Neduba diabolica* | 25 | 28 | (Warchałowska-Śliwa 1998) |
| Tettigoniidae | *Neduba macneilli* | 25 | 28 | (Warchałowska-Śliwa 1998) |
| Tettigoniidae | *Neduba diminitiva* | 23 | 25 | (Warchałowska-Śliwa 1998) |
| Tettigoniidae | *Neduba ovata armiger* | 23 | 23 | (Warchałowska-Śliwa 1998) |
| Tettigoniidae | *Neduba sp.* | 22 | 27 | (Warchałowska-Śliwa 1998) |
| Tettigoniidae | *Oligodectoides tindalei* | 29 | 29 | (Warchałowska-Śliwa 1998) |
| Tettigoniidae | *Oligodectes mallee* | 29 | 29 | (Warchałowska-Śliwa 1998) |
| Tettigoniidae | *Oligodectes pallens* | 29 | 29 | (Warchałowska-Śliwa 1998) |
| Tettigoniidae | *Oligodectes longicerus* | 21 | 21 | (Warchałowska-Śliwa 1998) |
| Tettigoniidae | *Onconotus laxmmani* | 25 | 27 | (Warchałowska-Śliwa 1998) |
| Tettigoniidae | *Paratlantlica sp.* | 25 | 27 | (Warchałowska-Śliwa 1998) |
| Tettigoniidae | *Paratlanticus ussuriensis* | 27 | 29 | (Warchałowska-Śliwa 1998) |
| Tettigoniidae | *Parapholidoptera naxia* | 31 | 31 | (Warchałowska-Śliwa 1998) |
| Tettigoniidae | *Parapholidoptera signata* | 31 | 31 | (Warchałowska-Śliwa et al. 2005) |
| Tettigoniidae | *Pholidoptera aptera aptera* | 29 | 31 | (Warchałowska-Śliwa 1998) |
| Tettigoniidae | *Pholidoptera aptera karnyi* | 29 | 31 | (Warchałowska-Śliwa 1998) |
| Tettigoniidae | *Pholidoptera frivaldsky* | 31 | 31 | (Warchałowska-Śliwa 1998) |
| Tettigoniidae | *Pholidoptera giseoaptera* | 31 | 31 | (Warchałowska-Śliwa 1998) |
| Tettigoniidae | *Pholidoptera macedonica* | 29 | 31 | (Warchałowska-Śliwa 1998) |
| Tettigoniidae | *Plagiostira gillettei* | 25 | 30 | (Warchałowska-Śliwa 1998) |

| | | | | |
|---|---|---|---|---|
| Tettigoniidae | *Platycleis affinis* | 31 | 31 | (Warchałowska-Śliwa 1998) |
| Tettigoniidae | *Platycleis albupunctata hispanica* | 31 | 31 | (Warchałowska-Śliwa 1998) |
| Tettigoniidae | *Platycleis denticulata* | 31 | 31 | (Warchałowska-Śliwa 1998) |
| Tettigoniidae | *Platycleis falx laticauda* | 31 | 31 | (Warchałowska-Śliwa 1998) |
| Tettigoniidae | *Platycleis grisea* | 31 | 31 | (Warchałowska-Śliwa 1998) |
| Tettigoniidae | *Platycleis intermedia* | 31 | 31 | (Warchałowska-Śliwa 1998) |
| Tettigoniidae | *Platycleis pamirica* | 31 | 31 | (Warchałowska-Śliwa 1998) |
| Tettigoniidae | *Platycleis tenuis* | 31 | 31 | (Warchałowska-Śliwa et al. 2005) |
| Tettigoniidae | *Platycleis tessellata* | 31 | 31 | (Warchałowska-Śliwa 1998) |
| Tettigoniidae | *Platycleis vittata* | 31 | 31 | (Warchałowska-Śliwa 1998) |
| Tettigoniidae | *Platydecticus angustifrons* | 37 | 37 | (Warchałowska-Śliwa 1998) |
| Tettigoniidae | *Psorodonotus illyricus macedonicus* | 33 | 33 | (Warchałowska-Śliwa et al. 2005) |
| Tettigoniidae | *Psorodonotus specularis* | 33 | 33 | (Warchałowska-Śliwa 1998) |
| Tettigoniidae | *Pterolepis ferdinandi* | 25 | 27 | (Warchałowska-Śliwa et al. 2005) |
| Tettigoniidae | *Pterolepis germanica* | 25 | 27 | (Warchałowska-Śliwa et al. 2005) |
| Tettigoniidae | *Pterolepis insularis* | 25 | 27 | (Warchałowska-Śliwa et al. 2005) |
| Tettigoniidae | *Pterolepis sp. nova* | 25 | 27 | (Warchałowska-Śliwa et al. 2005) |
| Tettigoniidae | *Pterolepis edentata* | 25 | 27 | (Warchałowska-Śliwa et al. 2005) |
| Tettigoniidae | *Rhachidorus blackdownensis* | 31 | 31 | (Warchałowska-Śliwa 1998) |
| Tettigoniidae | *Rhachidorus longipennis* | 31 | 31 | (Warchałowska-Śliwa 1998) |
| Tettigoniidae | *Rhachidorus semoni* | 31 | 31 | (Warchałowska-Śliwa 1998) |
| Tettigoniidae | *Steiroxys strepens* | 29 | 31 | (Warchałowska-Śliwa 1998) |
| Tettigoniidae | *Steiroxys trillineatus* | 29 | 29 | (Warchałowska-Śliwa 1998) |
| Tettigoniidae | *Steiroxys sp.* | 29 | 31 | (Warchałowska-Śliwa 1998) |
| Tettigoniidae | *Tadzhikia pavlovskii* | 27 | 31 | (Warchałowska-Śliwa 1998) |
| Tettigoniidae | *Tettigonia orientalis* | 35 | 35 | (Warchałowska-Śliwa 1998) |
| Tettigoniidae | *Tettigonia orientalis orientalis* | 33 | 33 | (Warchałowska-Śliwa 1998) |
| Tettigoniidae | *Tettigonia cantans* | 29 | 33 | (Warchałowska-Śliwa 1998) |
| Tettigoniidae | *Tettigonia caudata* | 29 | 32 | (Warchałowska-Śliwa 1998) |

| | | | | |
|---|---|---|---|---|
| Tettigoniidae | *Tettigonia viridissima* | 29 | 32 | (Warchałowska-Śliwa 1998) |
| Tettigoniidae | *Tettigonia sp.* | 29 | 31 | (Warchałowska-Śliwa 1998) |
| Tettigoniidae | *Tettigonia ussuriana* | 29 | 32 | (Warchałowska-Śliwa 1998) |
| Tettigoniidae | *Tettigonia dolichopoda maritima* | 29 | 32 | (Warchałowska-Sliwa et al. 2002) |
| Tettigoniidae | *Uvarovina daurica* | 31 | 31 | (Warchałowska-Śliwa 1998) |
| Tettigoniidae | *Xederra barbarae* | 33 | 33 | (Warchałowska-Śliwa 1998) |
| Tettigoniidae | *Zacycloptera atripennis* | 23 | 30 | (Warchałowska-Śliwa 1998) |
| Tettigoniidae | *Kawanaphila goolwa* | 35 | 35 | (Warchałowska-Śliwa 1998) |
| Tettigoniidae | *Kawanaphila iyouta* | 35 | 35 | (Warchałowska-Śliwa 1998) |
| Tettigoniidae | *Kawanaphila lexceni* | 35 | 35 | (Warchałowska-Śliwa 1998) |
| Tettigoniidae | *Kawanaphila mirla* | 35 | 35 | (Warchałowska-Śliwa 1998) |
| Tettigoniidae | *Kawanaphila nartee* | 35 | 35 | (Warchałowska-Śliwa 1998) |
| Tettigoniidae | *Kawanaphila triodiae* | 35 | 35 | (Warchałowska-Śliwa 1998) |
| Tettigoniidae | *Kawanaphila ungarunya* | 35 | 35 | (Warchałowska-Śliwa 1998) |
| Tettigoniidae | *Kawanaphila yarraga* | 35 | 35 | (Warchałowska-Śliwa 1998) |
| Tettigoniidae | *Windbalea warooa* | 35 | 35 | (Warchałowska-Śliwa 1998) |
| Tettigoniidae | *Windbalea viride* | 35 | 35 | (Warchałowska-Śliwa 1998) |
| Tettigoniidae | *Anthophiloptera dryas* | 35 | 35 | (Warchałowska-Śliwa 1998) |
| Tettigoniidae | *Phasmodes jaeba* | 31 | 33 | (Warchałowska-Śliwa 1998) |
| Tettigoniidae | *Phasmodes nungeroo* | 31 | 33 | (Warchałowska-Śliwa 1998) |
| Tettigoniidae | *Phasmodes ranatriformis* | 29 | 36 | (Warchałowska-Śliwa 1998) |
| Tettigoniidae | *Zaprochilus australis* | 31 | 31 | (Warchałowska-Śliwa 1998) |
| Tettigoniidae | *Zaprochilus ninae* | 31 | 31 | (Warchałowska-Śliwa 1998) |
| Tettigoniidae | *Zaprochilus mongabarra* | 31 | 31 | (Warchałowska-Śliwa 1998) |
| Rhaphidophoridae | *Diestrammena japonica* | 57 | 58 | (Mesa et al. 1968) |
| Rhaphidophoridae | *Diestrammena unicolor unicolor* | 29 | 29 | (Warchałowska-Śliwa and Kostia 1996) |
| Rhaphidophoridae | *Diestrammena Tachycines asynamorus* | 57 | 58 | (Mesa 1965) |
| Rhaphidophoridae | *Tachycines coreanus* | 49 | 49 | (Warchałowska-Śliwa and Kostia 1996) |
| Rhaphidophoridae | *Paratachycines (Hemitachycines) boldyrevi* | 47 | 47 | (Warchałowska-Śliwa and Kostia 1996) |

| | | | | |
|---|---|---|---|---|
| Rhaphidophoridae | *Ceuthophilus maculatus* | 37 | 37 | (Mesa 1965) |
| Rhaphidophoridae | *Ceuthophilus maculatus* | 39 | 39 | (Mesa 1965) |
| Rhaphidophoridae | *Ceuthophilus sp.* | 37 | 37 | (Mesa 1965) |
| Rhaphidophoridae | *Dolichopoda schiavazzii* | 32 | 64 | (Di Russo et al. 1994) |
| Rhaphidophoridae | *Dolichopoda linderi* | 28 | 56 | (Mesa et al. 1968) |
| Rhaphidophoridae | *Dolichopoda baccetti* | 31 | 62 | (Mesa 1965) |
| Rhaphidophoridae | *Dolichopoda laetitia* | 31 | 62 | (Mesa 1965) |
| Rhaphidophoridae | *Dolichopoda geniculata* | 31 | 62 | (Mesa 1965) |
| Rhaphidophoridae | *Dolichopoda ligustica* | 31 | 62 | (Mesa 1965) |
| Rhaphidophoridae | *Australotettix montanus* | 45 | 45 | (Mesa et al. 1968) |
| Rhaphidophoridae | *Cavernotettix buchanensis* | 39 | 39 | (Mesa et al. 1968) |
| Rhaphidophoridae | *Cavernotettix montanus* | 43 | 43 | (Mesa et al. 1968) |
| Rhaphidophoridae | *Cavernotettix wyanbenensis* | 34 | 34 | (Mesa et al. 1968) |
| Rhaphidophoridae | *Cavernotettix sp. nov. 1* | 44 | 46 | (Mesa et al. 1968) |
| Rhaphidophoridae | *Cavernotettix sp. nov. 2* | 43 | 47 | (Mesa et al. 1968) |
| Rhaphidophoridae | *Heteromallus spina* | 45 | 50 | (Mesa 1965) |
| Rhaphidophoridae | *Heteromallus gracilipes* | 45 | 50 | (Mesa 1965) |
| Rhaphidophoridae | *Heteromallu pectinipes* | 45 | 50 | (Mesa 1965) |
| Rhaphidophoridae | *Heteromallu spinifer* | 45 | 50 | (Mesa 1965) |
| Rhaphidophoridae | *Microphatus cavernicola* | 45 | 46 | (Mesa et al. 1968) |
| Rhaphidophoridae | *Microphatus tasmaniensis* | 45 | 46 | (Mesa et al. 1968) |
| Rhaphidophoridae | *Troglophilus neglectus* | 17 | 18 | (Mesa et al. 1968) |
| Rhaphidophoridae | *Troglophilus cavicola* | 21 | 21 | (Mesa 1965) |
| Anostostomatidae | *Apteranabropsis tonkinensis* | 19 | 30 | (Warchałowska-Śliwa et al. 1999) |
| Anostostomatidae | *Australostoma opacum* | 21 | 28 | (Warchałowska-Śliwa et al. 1999) |
| Anostostomatidae | *Deinacrida connectens* | 17 | 26 | (Morgan-Richards and Gibbs 2001) |
| Anostostomatidae | *Deinacrida connectens* | 19 | 28 | (Morgan-Richards and Gibbs 2001) |
| Anostostomatidae | *Deinacrida connectens* | 21 | 30 | (Morgan-Richards and Gibbs 2001) |
| Anostostomatidae | *Deinacrida heteracantha* | 21 | 28 | (Morgan-Richards and Gibbs 2001) |

| | | | | |
|---|---|---|---|---|
| Anostostomatidae | *Deinacrida fallai* | 21 | 28 | (Morgan-Richards and Gibbs 2001) |
| Anostostomatidae | *Deinacrida mahoenui* | 21 | 28 | (Morgan-Richards and Gibbs 2001) |
| Anostostomatidae | *Deinacrida elegans* | 27 | 28 | (Morgan-Richards and Gibbs 2001) |
| Anostostomatidae | *Deinacrida rugosa* | 29 | 30 | (Morgan-Richards and Gibbs 2001) |
| Anostostomatidae | *Deinacrida parva* | 29 | 30 | (Morgan-Richards and Gibbs 2001) |
| Anostostomatidae | *Deinacrida pluvialis* | 23 | 30 | (Morgan-Richards and Gibbs 2001) |
| Anostostomatidae | *Deinacrida talpa* | 23 | 30 | (Morgan-Richards and Gibbs 2001) |
| Anostostomatidae | *Deinacrida tibiospina* | 25 | 30 | (Morgan-Richards and Gibbs 2001) |
| Anostostomatidae | *Hemideina broughi* | 25 | 30 | (Morgan-Richards and Gibbs 2001) |
| Anostostomatidae | *Hemideina femorata* | 25 | 30 | (Morgan-Richards and Gibbs 2001) |
| Anostostomatidae | *Hemideina maori* | 25 | 30 | (Morgan-Richards and Gibbs 2001) |
| Anostostomatidae | *Hemideina ricta* | 25 | 30 | (Morgan-Richards and Gibbs 2001) |
| Anostostomatidae | *Hemideina thoracica* | 11 | 22 | (Morgan-Richards and Gibbs 2001) |
| Anostostomatidae | *Hemideina thoracica* | 13 | 22 | (Morgan-Richards and Gibbs 2001) |
| Anostostomatidae | *Hemideina thoracica* | 15 | 24 | (Morgan-Richards and Gibbs 2001) |
| Anostostomatidae | *Hemideina thoracica* | 17 | 26 | (Morgan-Richards and Gibbs 2001) |
| Anostostomatidae | *Hemideina thoracica* | 19 | 28 | (Morgan-Richards and Gibbs 2001) |
| Anostostomatidae | *Hemideina thoracica* | 23 | 30 | (Morgan-Richards and Gibbs 2001) |
| Anostostomatidae | *Hemideina trewicki* | 17 | 32 | (Morgan-Richards and Gibbs 2001) |
| Anostostomatidae | *Hemideina crassidens* | 13 | 24 | (Morgan-Richards and Gibbs 2001) |
| Anostostomatidae | *Hemideina crassidens* | 19 | 28 | (Morgan-Richards and Gibbs 2001) |
| Gryllacrididae | *Gryllacris signifera* | 11 | 11 | (Mesa 1965) |
| Gryllacrididae | *Nippancistroger testaceus* | 17 | 17 | (Mesa 1965) |
| Gryllotalpidae | *Gryllotalpa africana* | 23 | 31 | (White 1973) |
| Gryllotalpidae | *Gryllotalpa borealis* | 23 | 31 | (White 1973) |
| Gryllotalpidae | *Gryllotalpa parva* | 23 | 31 | (White 1973) |
| Gryllotalpidae | *Gryllotalpa fossor* | 23 | 31 | (Rao and Arora 1979) |
| Gryllotalpidae | *Gryllotalpa himalayana* | 23 | 31 | (White 1973) |
| Gryllotalpidae | *Gryllotalpa vulgaris* | 23 | 31 | (Honda 1926) |

| | | | | |
|---|---|---|---|---|
| Gryllotalpidae | *Gryllotalpa gryllotalpa* | 23 | 31 | (White 1973) |
| Gryllotalpidae | *Neocurtilla (Gryllotalpa) hexadactyla* | 23 | 31 | (White 1973) |
| Gryllotalpidae | *Gryllotalpa septemdecimchromosomica* | 17 | 31 | (White 1973) |
| Gryllotalpidae | *Gryllotalpa marismortui 1* | 23 | 31 | (Broza et al. 1998) |
| Gryllotalpidae | *Gryllotalpa marismortui 2* | 19 | 31 | (Broza et al. 1998) |
| Gryllotalpidae | *Gryllotalpa cossyrensis 1* | 23 | 31 | (Broza et al. 1998) |
| Gryllotalpidae | *Gryllotalpa cossyrensis 2* | 20 | 31 | (Broza et al. 1998) |
| Gryllotalpidae | *Gryllotalpa cossyrensis 3* | 18 | 31 | (Broza et al. 1998) |
| Gryllotalpidae | *Gryllotalpa cossyrensis 4* | 16 | 31 | (Broza et al. 1998) |
| Gryllotalpidae | *Gryllotalpa cossyrensis 5* | 15 | 31 | (Broza et al. 1998) |
| Gryllotalpidae | *Gryllotalpa. tali* | 19 | 31 | (Broza et al. 1998) |
| Gryllotalpidae | *Gryllotalpa krimbasi* | 19 | 31 | (Broza et al. 1998) |
| Gryllotalpidae | *Scapteriscus tetradactylus* | 23 | 31 | (White 1973) |
| Stenopelmatidae | *Stenoplematus ssp.* | 47 | 47 | (Mesa 1965) |
| Stenopelmatidae | *Lutosa brasiliensis* | 15 | 15 | (Warchałowska-Śliwa et al. 1999) |
| Cylindrachetidae | *Cylinbdracheta psammofhila* | 15 | 18 | (John and Rentz 1987) |
| Cylindrachetidae | *Cylindroryctes spegazzinii* | 13 | 17 | (Mesa 1977) |
| Tridactylidae | *Tridactylus variegatus* | 11 | 21 | (John and Rentz 1987) |
| Tridactylidae | *Tridactylus mutus mutus* | 13 | 26 | (John and Rentz 1987) |
| Tridactylidae | *Tridactylus japonicus* | 13 | 23 | (John and Rentz 1987) |
| Ripipterygidae | *Rhipipteryx nolata* | 15 | 29 | (John and Rentz 1987) |
| Tetrigidae | *Formasatettix robustus* | 13 | 13 | (Warchałowska-Śliwa et al. 2003) |
| Tetrigidae | *Paratettix meridionalis* | 13 | 13 | (Warchałowska-Śliwa et al. 2003) |
| Tetrigidae | *Tetrix bolivari* | 13 | 13 | (Warchałowska-Śliwa et al. 2003) |
| Tetrigidae | *Tetrix japonica* | 13 | 13 | (Warchałowska-Śliwa et al. 2003) |
| Tetrigidae | *Tetrix simulans* | 13 | 13 | (Warchałowska-Śliwa et al. 2003) |
| Tetrigidae | *Tetrix subulata* | 13 | 13 | (Warchałowska-Śliwa et al. 2003) |
| Tetrigidae | *Tetrix tenuicornis* | 13 | 13 | (Warchałowska-Śliwa et al. 2003) |
| Tetrigidae | *Tetrix undulata* | 13 | 13 | (Warchałowska-Śliwa et al. 2003) |

| | | | | |
|---|---|---|---|---|
| Tetrigidae | *Tetrix wagai* | 13 | 13 | (Warchałowska-Śliwa et al. 2003) |
| Tetrigidae | *Uvarovittetix depressus* | 13 | 13 | (Warchałowska-Śliwa et al. 2003) |
| Proscopiidae | *Stiphra robusta* | 19 | 19 | (Moura et al. 1996) |
| Proscopiidae | *Tetanorhynchus silvai* | 19 | 19 | (Moura et al. 1996) |
| Proscopiidae | *Tetanorhynchus mendesi* | 17 | 21 | (Moura et al. 1996) |
| Proscopiidae | *Scleratoscopia protopeirae* | 19 | 24 | (Moura et al. 1996) |
| Proscopiidae | *Scleratoscopia spinosa* | 19 | 24 | (Moura et al. 1996) |
| Proscopiidae | *Cephalocoema zilkari* | 17 | 21 | (Moura et al. 1996) |
| Proscopiidae | *Anchocoema sp 1* | 15 | 19 | (Moura et al. 1996) |
| Proscopiidae | *Anchocoema sp 2* | 15 | 19 | (Moura et al. 1996) |
| Proscopiidae | *Hybusa armaticolis* | 17 | 21 | (Moura et al. 1996) |
| Epistacidae | *Seyrigella notabilis* | 25 | 25 | (White 1970) |
| Epistacidae | *Heteromastax appendiculata* | 25 | 25 | (White 1970) |
| Eumastacidae | *Xenomastax wintreberti* | 19 | 19 | (White 1970) |
| Eumastacidae | *Apteropeoedes pygmaes* | 21 | 23 | (White 1970) |
| Eumastacidae | *Apteropeoedes elegans* | 21 | 21 | (White 1970) |
| Eumastacidae | *Apteropeoedes wintreberti* | 21 | 21 | (White 1970) |
| Eumastacidae | *Apteropeoedes rostratus* | 21 | 21 | (White 1970) |
| Eumastacidae | *Tetefortina wintreberti* | 19 | 21 | (White 1970) |
| Eumastacidae | *Tetefortina curta* | 19 | 21 | (White 1970) |
| Eumastacidae | *Micromastax teteforti rectifrons* | 21 | 21 | (White 1970) |
| Eumastacidae | *Micromastax teteforti cavifrons* | 21 | 21 | (White 1970) |
| Eumastacidae | *Lavanonia thalassina* | 21 | 21 | (White 1970) |
| Eumastacidae | *Lavanonia itampolae* | 21 | 21 | (White 1970) |
| Eumastacidae | *Wintrebertia angulata* | 21 | 21 | (White 1970) |
| Eumastacidae | *Wintrebertia arcuata* | 21 | 21 | (White 1970) |
| Eumastacidae | *Wintrebertia teteforti andranovatae* | 21 | 21 | (White 1970) |
| Eumastacidae | *Wintrebertia lavanoniae* | 21 | 21 | (White 1970) |
| Eumastacidae | *Wintrebertia callosa* | 21 | 21 | (White 1970) |

| | | | | |
|---|---|---|---|---|
| Eumastacidae | *Wintrebertia crassipes* | 21 | 21 | (White 1970) |
| Eumastacidae | *Parawintrebertia gigantea* | 21 | 21 | (White 1970) |
| Eumastacidae | *Parawintrebertia pauliani pauliani* | 21 | 21 | (White 1970) |
| Eumastacidae | *Exophatalmomastax malzyi* | 21 | 21 | (White 1970) |
| Eumastacidae | *Parapisactuas carinatus* | 19 | 28 | (White 1970) |
| Eumastacidae | *Paramastax rosenbergi* | 19 | 28 | (White 1970) |
| Eumastacidae | *Eumastax salazari* | 21 | 21 | (White 1970) |
| Morabidae | *Vamdiemenella viatica 19* | 19 | 22 | (White 1978) |
| Morabidae | *Vamdiemenella viatica 17* | 17 | 22 | (White 1978) |
| Morabidae | *P24X0* | 17 | 21 | (White 1978) |
| Morabidae | *P24XY* | 16 | 25 | (White 1978) |
| Morabidae | *P25X0* | 19 | 21 | (White 1978) |
| Morabidae | *P25XY* | 18 | 21 | (White 1978) |
| Morabidae | *P45bX0* | 19 | 21 | (White 1978) |
| Morabidae | *P45bXY* | 18 | 21 | (White 1978) |
| Morabidae | *Keyacris scurra 15* | 15 | 15 | (White 1978) |
| Morabidae | *Keyacris scurra 17* | 17 | 17 | (White 1978) |
| Morabidae | *Warramba virgo* | 15 | 22 | (Webb and Westerman 1978) |
| Morabidae | *Warramba picta* | 17 | 21 | (Webb and Westerman 1978) |
| Morabidae | *Warramba P169* | 16 | 22 | (Schweizer, et al. 1983) |
| Morabidae | *Warramba P196* | 14 | 22 | (Schweizer, et al. 1983) |
| Pyrgomorphidae | *Omura congrua* | 19 | 19 | (Mesa et al. 1982) |
| Pyrgomorphidae | *Algete brunneri* | 19 | 19 | (Anjos et al. 2015) |
| Pyrgomorphidae | *Pyrgomorpha conica* | 19 | 19 | Anjos et al. 2015) |
| Pyrgomorphidae | *Pyrgomorpha rugosa* | 11 | 19 | (Fossey et al. 1989) |
| Pyrgomorphidae | *Pyrgomorpha granulata* | 13 | 19 | (Fossey et al. 1989) |
| Pyrgomorphidae | *Pyrgomorpha sp* | 19 | 19 | (Fossey et al. 1989) |
| Pyrgomorphidae | *Pyrgomorpha guentheri* | 19 | 19 | (Buleu et al. 2017) |
| Pyrgomorphidae | *Zonocerus elegans* | 19 | 19 | (Buleu et al. 2017) |

| | | | | |
|---|---|---|---|---|
| Pyrgomorphidae | *Atractomorpha lata* | 19 | 19 | (Buleu et al. 2017) |
| Pamphagidae | *Acinipe calabra* | 19 | 19 | (Warchałowska-Śliwa et al. 1994) |
| Pamphagidae | *Haplotropis brunneriana* | 19 | 19 | (Bugrov and Warchałowska-Śliwa 1997) |
| Pamphagidae | *Nocaracris cyanipes* | 18 | 19 | (Bugrov and Warchałowska-Śliwa 1997; Castillo et al. 2010) |
| Pamphagidae | *Pamphagus cristatus* | 19 | 19 | (Warchałowska-Śliwa et al. 1994) |
| Pamphagidae | *Pamphagus marmoratus* | 19 | 19 | (Warchałowska-Śliwa et al. 1994) |
| Pamphagidae | *Paranocaracris bulgaricus* | 18 | 19 | (Bugrov and Grozev A 1998) |
| Pamphagidae | *Paranocarodes straubei* | 18 | 19 | (Bugrov and Grozev A 1998) |
| Pamphagidae | *Paranocarodes chopardi* | 18 | 19 | (Bugrov and Grozev A 1998) |
| Pamphagidae | *Paranothrotes opacus* | 18 | 19 | (Bugrov et al. 2016) |
| Pamphagidae | *Paranothrotes rubripes* | 18 | 19 | (Bugrov et al. 2016) |
| Pamphagidae | *Saxetania cultricollis* | 18 | 19 | (Bugrov and Warchałowska-Śliwa 1997; Castillo et al. 2010) |
| Pamphagidae | *Asiotmethis heptapotamicus* | 18 | 19 | (Bugrov and Grozev A 1998; Castillo et al. 2010) |
| Pamphagidae | *Asiotmethis limbatus* | 18 | 19 | (Bugrov and Grozev A 1998) |
| Pamphagidae | *Asiotmethis turritus* | 18 | 19 | (Bugrov et al. 2016) |
| Pamphagidae | *Asiotmethis zacharjini* | 18 | 19 | (Castillo et al. 2010) |
| Pamphagidae | *Atrichotmethis semenovi* | 18 | 19 | (Bugrov and Warchałowska-Śliwa 1997) |
| Pamphagidae | *Eromopeza festiva* | 18 | 19 | (Bugrov et al. 2016) |
| Pamphagidae | *Melanotmethis fuscipennis* | 19 | 28 | (Bugrov and Warchałowska-Śliwa 1997) |
| Pamphagidae | *Strumiger desertorum* | 19 | 19 | (Bugrov and Warchałowska-Śliwa 1997) |
| Pamphagidae | *Thrinchus arenosus* | 19 | 19 | (Bugrov and Warchałowska-Śliwa 1997) |
| Pamphagidae | *Eumigus punctatus* | 19 | 19 | (Camacho et al. 1981) |
| Pamphagidae | *Eumigus monticulus* | 19 | 19 | (Camacho et al. 1981) |
| Pamphagidae | *Eumigus cucullatus* | 19 | 19 | (Camacho et al. 1981) |
| Lentulidae | *Karruacris browni* | 20 | 23 | (White 1967) |
| Lentulidae | *Karruacris browni* | 19 | 23 | (White 1967) |
| Lentulidae | *Karruia paradoxa* | 23 | 23 | (White 1967) |
| Lentulidae | *Lentula callani* | 23 | 23 | (White 1967) |
| Lentulidae | *Mecostibus cf. nyassae* | 23 | 23 | (White 1967) |

| | | | | |
|---|---|---|---|---|
| Lentulidae | *Shelfordites nanus* | 21 | 21 | (White 1967) |
| Lentulidae | *Syrgus sp.* | 23 | 23 | (White 1967) |
| Lentulidae | *Paralentula marcida* | 23 | 23 | (White 1967) |
| Lentulidae | *Paralentula prasinata* | 23 | 23 | (White 1967) |
| Lentulidae | *Lentula n. sp.* | 23 | 23 | (White 1967) |
| Lentulidae | *Basutacris n. sp. cf. minuta* | 23 | 23 | (White 1967) |
| Lentulidae | *Eremidium denticercus* | 23 | 23 | (White 1967) |
| Tristiridae | *Atamacris diminuta* | 20 | 23 | (Castillo et al. 2010) |
| Tristiridae | *Bufonacris sp* | 19 | 23 | (Mesa et al. 1982) |
| Tristiridae | *Elasmoderus rabiosus* | 23 | 23 | (Mesa et al. 1982) |
| Tristiridae | *Elysiacris angusticollis* | 21 | 21 | (Mesa et al. 1982) |
| Tristiridae | *Moluchacris cinerascens* | 21 | 23 | (Mesa et al. 1982) |
| Tristiridae | *Peplacris recutita* | 21 | 23 | (Mesa et al. 1982) |
| Tristiridae | *Tropidostethus bicarinatus* | 21 | 21 | (Mesa et al. 1982) |
| Tristiridae | *Illapelia penai* | 23 | 23 | (Mesa et al. 1982) |
| Acrididae | *Catantops humilis* | 23 | 23 | (Castillo et al. 2010) |
| Acrididae | *Catantops humilis* | 22 | 23 | (Castillo et al. 2010) |
| Acrididae | *Aleuas gracilis* | 20 | 23 | (Castillo et al. 2010) |
| Acrididae | *Aleuas lineatus* | 20 | 23 | (Castillo et al. 2010) |
| Acrididae | *Aleuas vitticolis* | 20 | 23 | (Castillo et al. 2010) |
| Acrididae | *Aleuas paranensis* | 20 | 23 | (Castillo et al. 2010) |
| Acrididae | *Aleuas paraguayensis* | 22 | 23 | (Castillo et al. 2010) |
| Acrididae | *Aleuas albinae* | 20 | 23 | (Castillo et al. 2010) |
| Acrididae | *Zygoclistron falconicum* | 20 | 23 | (Castillo et al. 2010) |
| Acrididae | *Zygoclistron nasicum* | 20 | 23 | (Castillo et al. 2010) |
| Acrididae | *Zygoclistron trachysticum* | 20 | 23 | (Castillo et al. 2010) |
| Acrididae | *Eyprepocnemis unicolor* | 23 | 23 | (Bugrov et al. 1999) |
| Acrididae | *Eyprepocnemis plorans* | 23 | 23 | (Bugrov et al. 1999) |
| Acrididae | *Heterachris adspersa* | 23 | 23 | (Bugrov et al. 1999) |

| | | | | |
|---|---|---|---|---|
| Acrididae | *Heterachris pulcher* | 22 | 23 | (Castillo et al. 2010) |
| Acrididae | *shirakiacris shirakii* | 23 | 23 | (Bugrov et al. 1999) |
| Acrididae | *Thisoicetrinus pterostichus* | 23 | 23 | (Bugrov et al. 1999) |
| Acrididae | *Mermiria bivittata* | 22 | 23 | (Castillo et al. 2010) |
| Acrididae | *Mermiria intertexta* | 22 | 23 | (Castillo et al. 2010) |
| Acrididae | *Mermiria maculipennis* | 22 | 23 | (Castillo et al. 2010) |
| Acrididae | *Scyllinula humilis* | 22 | 22 | (Castillo et al. 2010) |
| Acrididae | *Sinipta dalmani* | 23 | 23 | (Castillo et al. 2010) |
| Acrididae | *Sinipta dalmani* | 22 | 22 | (Castillo et al. 2010) |
| Acrididae | *Stenobothrus rubicundus* | 22 | 23 | (Castillo et al. 2010) |
| Acrididae | *Anapodisma miramae* | 22 | 23 | (Bugrov et al. 1994) |
| Acrididae | *Atrachelacris unicolor* | 22 | 23 | (Castillo et al. 2010) |
| Acrididae | *Atrachelacris olivaceus* | 22 | 23 | (Castillo et al. 2010) |
| Acrididae | *Baeacris punctulatus* | 23 | 23 | (Castillo et al. 2010) |
| Acrididae | *Baeacris punctulatus* | 22 | 23 | (Castillo et al. 2010) |
| Acrididae | *Boliviacris noroestensis* | 20 | 21 | (Castillo et al. 2010) |
| Acrididae | *Dichroplus maculipennis* | 22 | 23 | (Castillo et al. 2010) |
| Acrididae | *Dichroplus obscurus* | 18 | 23 | (Castillo et al. 2010) |
| Acrididae | *Dichroplus porteri* | 22 | 23 | (Castillo et al. 2010) |
| Acrididae | *Dichroplus silveiraguidoi* | 8 | 13 | (Castillo et al. 2010) |
| Acrididae | *Dichroplus vittatus* | 18 | 23 | (Castillo et al. 2010) |
| Acrididae | *Dichroplus vittigerum* | 18 | 23 | (Castillo et al. 2010) |
| Acrididae | *Dichromatos lilloanus* | 21 | 23 | (Palacios-Gimenez et al. 2013) |
| Acrididae | *Dichromatos schrottkyi* | 21 | 23 | (Palacios-Gimenez et al. 2013) |
| Acrididae | *Dichromatos corupa* | 21 | 23 | (Castillo et al. 2010) |
| Acrididae | *Dichromatos montanus* | 21 | 23 | (Castillo et al. 2010) |
| Acrididae | *Fruhstorferiola okinawaensis* | 21 | 21 | (Bugrov et al. 2000) |
| Acrididae | *Hesperotettix speciosus* | 22 | 22 | (Castillo et al. 2010) |
| Acrididae | *Hesperotettix viridis* | 22 | 22 | (Castillo et al. 2010) |

| | | | | |
|---|---|---|---|---|
| Acrididae | *Hesperotettix pratensis* | 22 | 22 | (Castillo et al. 2010) |
| Acrididae | *Hypochlora alba* | 22 | 22 | (Castillo et al. 2010) |
| Acrididae | *Eirenephilus longipennis* | 23 | 23 | (Castillo et al. 2010) |
| Acrididae | *Eurotettix minor* | 22 | 23 | (Palacios-Gimenez et al. 2013) |
| Acrididae | *Leiotettix flavipes* | 22 | 23 | (Castillo et al. 2010) |
| Acrididae | *Leiotettix politus* | 14 | 19 | (Castillo et al. 2010) |
| Acrididae | *Leiotettix politus* | 13 | 19 | (Mesa et al. 1982) |
| Acrididae | *Leiotettix pulcher* | 22 | 23 | (Castillo et al. 2010) |
| Acrididae | *Leiotettix sanguineus* | 23 | 23 | (Castillo et al. 2010) |
| Acrididae | *Leiotettix sanguineus* | 22 | 23 | (Castillo et al. 2010) |
| Acrididae | *Mariacris viridipes* | 20 | 19 | (Castillo et al. 2010) |
| Acrididae | *Melanoplus frigidus* | 23 | 23 | (Castillo et al. 2010) |
| Acrididae | *Miramella alpina* | 21 | 21 | (Castillo et al. 2010) |
| Acrididae | *Miramella solitaria* | 21 | 21 | (Castillo et al. 2010) |
| Acrididae | *Oedaleonotus enigma* | 22 | 23 | (Castillo et al. 2010) |
| Acrididae | *Paratylotropidia beutemulleri* | 22 | 23 | (Castillo et al. 2010) |
| Acrididae | *Paratylotropidia brunneri* | 19 | 23 | (Castillo et al. 2010) |
| Acrididae | *Paratylotropidia morsei* | 19 | 23 | (Castillo et al. 2010) |
| Acrididae | *Perixerus squamipennis* | 20 | 23 | (Castillo et al. 2010) |
| Acrididae | *Philocleon anomalus* | 22 | 23 | (Castillo et al. 2010) |
| Acrididae | *Podisma pedestris* | 23 | 23 | (Castillo et al. 2010) |
| Acrididae | *Podisma pedestris* | 22 | 23 | (John and Hewitt 1970) |
| Acrididae | *Podisma sapporensis* | 23 | 23 | (Bugrov et al. 2000) |
| Acrididae | *Podisma sapporensis* | 22 | 23 | (Bugrov et al. 2000) |
| Acrididae | *Podisma subastris* | 21 | 21 | (Bugrov et al. 2000) |
| Acrididae | *Parapodisma tenryuensis* | 21 | 21 | (Bugrov et al. 2000) |
| Acrididae | *Parapodisma yamato* | 21 | 21 | (Bugrov et al. 2000) |
| Acrididae | *Parapodisma mikado* | 21 | 21 | (Bugrov et al. 2000) |
| Acrididae | *Primnoa primnoides* | 23 | 23 | (Bugrov et al. 1994) |

| | | | | |
|---|---|---|---|---|
| Acrididae | *Primnoa primnoa* | 23 | 23 | (Bugrov et al. 1994) |
| Acrididae | *Primnoa litoralis* | 23 | 23 | (Bugrov et al. 1994) |
| Acrididae | *Primnoa plana* | 23 | 23 | (Bugrov et al. 1994) |
| Acrididae | *Podisma aberrans* | 23 | 23 | (Bugrov et al. 1994) |
| Acrididae | *Ronderosia bergi* | 22 | 23 | (Palacios-Gimenez et al. 2018) |
| Acrididae | *Ronderosia dubius* | 22 | 23 | (Castillo et al. 2010) |
| Acrididae | *Ronderosia forcipatus* | 22 | 23 | (Castillo et al. 2010) |
| Acrididae | *Ronderosia malloi* | 22 | 23 | (Castillo et al. 2010) |
| Acrididae | *Ronderosia ommexechoides* | 22 | 23 | (Castillo et al. 2010) |
| Acrididae | *Ronderosia paraguayensis* | 20 | 23 | (Castillo et al. 2010) |
| Acrididae | *Ronderosia piceomaculatus* | 22 | 23 | (Castillo et al. 2010) |
| Acrididae | *Ronderosia robustus* | 21 | 23 | (Castillo et al. 2010) |
| Acrididae | *Ronderosia cinctipes* | 21 | 23 | (Castillo et al. 2010) |
| Acrididae | *Scotussa daguerrei* | 21 | 23 | (Castillo et al. 2010) |
| Acrididae | *Scotussa delicatula* | 16 | 23 | (Castillo et al. 2010) |
| Acrididae | *Sinapodisma punctata* | 21 | 21 | (Bugrov et al. 2000) |
| Acrididae | *Zubovskya koeppeni* | 21 | 21 | (Bugrov et al. 1994) |
| Acrididae | *Tolgadia bivittata* | 22 | 23 | (Castillo et al. 2010) |
| Acrididae | *Tolgadia infirma* | 22 | 27 | (Castillo et al. 2010) |
| Acrididae | *Tolgadia sp.* | 22 | 23 | (Castillo et al. 2010) |
| Acrididae | *Zubovskya koreana* | 21 | 21 | Bugrov et al. 1994) |
| Acrididae | *Apacris sp* | 23 | 23 | (Mesa et al. 1982) |
| Acrididae | *Apacris rubrithorax* | 23 | 23 | (Mesa et al. 1982) |
| Acrididae | *Chlorus bolivianus* | 23 | 23 | (Mesa et al. 1982) |
| Acrididae | *Chlorus borrelli* | 23 | 23 | (Mesa et al. 1982) |
| Acrididae | *Chlorus vittatus* | 23 | 23 | (Palacios-Gimenez et al. 2013) |
| Acrididae | *Dichroplus alejomesai* | 23 | 23 | (Mesa et al. 1982) |
| Acrididae | *Dichroplus auriventris* | 23 | 23 | (Mesa et al. 1982) |
| Acrididae | *Dichroplus conspersus* | 23 | 23 | (Mesa et al. 1982) |

| | | | | |
|---|---|---|---|---|
| Acrididae | *Dichroplus democraticus* | 23 | 23 | (Mesa et al. 1982) |
| Acrididae | *Dichroplus elongatus* | 23 | 23 | (Mesa et al. 1982) |
| Acrididae | *Dichroplus exilis* | 23 | 23 | (Mesa et al. 1982) |
| Acrididae | *Dichroplus fuscus* | 19 | 23 | (Mesa et al. 1982) |
| Acrididae | *Dichroplus fuscus* | 20 | 23 | (Mesa et al. 1982) |
| Acrididae | *Dichroplus maculipennis* | 23 | 23 | (Mesa et al. 1982) |
| Acrididae | *Dichroplus mantiqueirae* | 23 | 23 | (Mesa et al. 1982) |
| Acrididae | *Dichroplus misionensis* | 23 | 23 | (Mesa et al. 1982) |
| Acrididae | *Dichroplus obsurus* | 18 | 22 | (Mesa et al. 1982) |
| Acrididae | *Dichroplus paraelongatus* | 23 | 23 | (Mesa et al. 1982) |
| Acrididae | *Dichroplus pratensis* | 19 | 19 | (Mesa et al. 1982) |
| Acrididae | *Dichroplus pratensis* | 18 | 19 | (Mesa et al. 1982) |
| Acrididae | *Dichroplus pratensis* | 17 | 19 | (Mesa et al. 1982) |
| Acrididae | *Dichroplus pratensis* | 16 | 19 | (Mesa et al. 1982) |
| Acrididae | *Dichroplus pratensis* | 15 | 19 | (Mesa et al. 1982) |
| Acrididae | *Dichroplus pratensis* | 14 | 19 | (Mesa et al. 1982) |
| Acrididae | *Dichroplus pratensis* | 13 | 19 | (Mesa et al. 1982) |
| Acrididae | *Dichroplus pseudopunctulatus* | 23 | 23 | (Mesa et al. 1982) |
| Acrididae | *Dichroplus robustus* | 21 | 23 | (Mesa et al. 1982) |
| Acrididae | *Dichroplus robustulus* | 23 | 23 | (Mesa et al. 1982) |
| Acrididae | *Dichroplus schulzi* | 23 | 23 | (Mesa et al. 1982) |
| Acrididae | *Leiotettix viridis* | 23 | 23 | (Mesa et al. 1982) |
| Acrididae | *Nahuelia rubriventris* | 23 | 23 | (Mesa et al. 1982) |
| Acrididae | *Neopedies brunneri* | 23 | 23 | (Mesa et al. 1982) |
| Acrididae | *Parascopas obesus* | 23 | 23 | (Mesa et al. 1982) |
| Acrididae | *Parascopas sanguineus* | 23 | 23 | (Mesa et al. 1982) |
| Acrididae | *Parascopas exertus* | 21 | 23 | (Mesa et al. 1982) |
| Acrididae | *Pedies andeaus* | 21 | 23 | (Mesa et al. 1982) |
| Acrididae | *Propedies bilobus* | 23 | 23 | (Mesa et al. 1982) |

| | | | | |
|---|---|---|---|---|
| Acrididae | *Propedies bipunctatus* | 23 | 23 | (Mesa et al. 1982) |
| Acrididae | *Propedies fusiformis* | 23 | 23 | (Mesa et al. 1982) |
| Acrididae | *Propedies olivaceus* | 23 | 23 | (Mesa et al. 1982) |
| Acrididae | *Propedies sanguineus* | 23 | 23 | (Mesa et al. 1982) |
| Acrididae | *Pseudoscopas nigrigena* | 23 | 23 | (Mesa et al. 1982) |
| Acrididae | *Pseudoscopas carbonelli* | 23 | 23 | (Mesa et al. 1982) |
| Acrididae | *Scotussa cliens* | 21 | 23 | (Mesa et al. 1982) |
| Acrididae | *Scotussa impudica* | 23 | 23 | (Mesa et al. 1982) |
| Acrididae | *Scotussa lemniscata* | 23 | 23 | (Mesa et al. 1982) |
| Acrididae | *Scotussa liebermanni* | 21 | 23 | (Mesa et al. 1982) |
| Acrididae | *Eucephalacris borellii* | 23 | 23 | (Mesa et al. 1982) |
| Acrididae | *Adimantus cubiceps* | 23 | 23 | (Mesa et al. 1982) |
| Acrididae | *Belosacris coccineipes* | 23 | 23 | (Mesa et al. 1982) |
| Acrididae | *Carbonellacris* | 23 | 23 | (Mesa et al. 1982) |
| Acrididae | *Leptysma dorsalis* | 23 | 23 | (Mesa et al. 1982) |
| Acrididae | *Leptysmina pallida* | 23 | 23 | (Mesa et al. 1982) |
| Acrididae | *Cornops frenatum* | 23 | 23 | (Mesa et al. 1982) |
| Acrididae | *Haroldgrantia lignosa* | 23 | 23 | (Mesa et al. 1982) |
| Acrididae | *Mastusia quadricarinata* | 23 | 23 | (Mesa et al. 1982) |
| Acrididae | *Oxyblepta sp* | 23 | 23 | (Mesa et al. 1982) |
| Acrididae | *Oxybleptella sagitta* | 23 | 23 | (Mesa et al. 1982) |
| Acrididae | *Stenopola boliviana* | 23 | 23 | (Mesa et al. 1982) |
| Acrididae | *Stenopola bohlsii* | 23 | 23 | (Mesa et al. 1982) |
| Acrididae | *Stenopola dorsalis* | 23 | 23 | (Mesa et al. 1982) |
| Acrididae | *Stenopola pallida* | 21 | 23 | (Mesa et al. 1982) |
| Acrididae | *Stenopola rubrifons rubrifons* | 23 | 23 | (Mesa et al. 1982) |
| Acrididae | *Tetrataenia surinama* | 19 | 23 | (Mesa et al. 1982) |
| Acrididae | *Paropaon laevifrons* | 23 | 23 | (Mesa et al. 1982) |
| Acrididae | *Paropaon pilosus tingomariae* | 23 | 23 | (Mesa et al. 1982) |

| | | | | |
|---|---|---|---|---|
| Acrididae | *Albretchia palpata* | 23 | 23 | (Mesa et al. 1982) |
| Acrididae | *Eulampiacris leucoptera* | 23 | 23 | (Mesa et al. 1982) |
| Acrididae | *Lamiacris migroguttata* | 23 | 23 | (Mesa et al. 1982) |
| Acrididae | *Machaeropeles rostratum* | 23 | 23 | (Mesa et al. 1982) |
| Acrididae | *Ommatolampis perspicillata* | 23 | 23 | (Mesa et al. 1982) |
| Acrididae | *Sitalces dorsalis* | 23 | 23 | (Mesa et al. 1982) |
| Acrididae | *Sitalces infuscatus* | 23 | 23 | (Mesa et al. 1982) |
| Acrididae | *Pycnosarcus atavus* | 23 | 23 | (Mesa et al. 1982) |
| Acrididae | *Abracris dilecta* | 23 | 23 | (Mesa et al. 1982) |
| Acrididae | *Abracris flavolineata* | 23 | 23 | (Bueno et al. 2013) |
| Acrididae | *Eujivarus fusiformis* | 21 | 23 | (Mesa et al. 1982) |
| Acrididae | *Eujivarus n. sp. A* | 21 | 23 | (Mesa et al. 1982) |
| Acrididae | *Eujivarus n. sp. B* | 21 | 23 | (Mesa et al. 1982) |
| Acrididae | *Eujivarus n sp. C* | 21 | 23 | (Mesa et al. 1982) |
| Acrididae | *Eujivarus vittatus* | 23 | 23 | (Mesa et al. 1982) |
| Acrididae | *Eusitalces vulneratus* | 23 | 23 | (Mesa et al. 1982) |
| Acrididae | *Jodacris ferrugineus ferrugineus* | 19 | 19 | (Mesa et al. 1982) |
| Acrididae | *Jodacris chapadensis* | 19 | 19 | (Mesa et al. 1982) |
| Acrididae | *Jodacris furcillata* | 19 | 19 | (Mesa et al. 1982) |
| Acrididae | *Omalotettix obliquum* | 21 | 23 | (Mesa et al. 1982) |
| Acrididae | *Osmilia flavolineata* | 23 | 23 | (Mesa et al. 1982) |
| Acrididae | *Psiloscirtus bolivianus* | 23 | 27 | (Mesa et al. 1982) |
| Acrididae | *Psiloscirtus olivaceus* | 23 | 27 | (Mesa et al. 1982) |
| Acrididae | *Psiloscirtus sp.* | 23 | 27 | (Mesa et al. 1982) |
| Acrididae | *Sitalces volxemi* | 19 | 23 | (Mesa et al. 1982) |
| Acrididae | *Xiphiola borellii* | 23 | 23 | (Mesa et al. 1982) |
| Acrididae | *Schistocerca cancellata* | 23 | 23 | (Mesa et al. 1982) |
| Acrididae | *Schistocerca flavofasciata* | 23 | 23 | (Mesa et al. 1982) |
| Acrididae | *Schistocerca pallens* | 23 | 23 | (Mesa et al. 1982) |

| | | | | |
|---|---|---|---|---|
| Acrididae | *Schistocerca paranensis* | 23 | 23 | (Mesa et al. 1982) |
| Acrididae | *Schistocerca gregaria* | 23 | 23 | (Palacios-Gimenez et al. 2020) |
| Acrididae | *Schistocerca serialis* | 23 | 23 | (Palacios-Gimenez et al. 2020) |
| Acrididae | *Schistocerca caribbeana* | 23 | 23 | (Palacios-Gimenez et al. 2020) |
| Acrididae | *Schistocerca cancellata* | 23 | 23 | (Palacios-Gimenez et al. 2020) |
| Acrididae | *Schistocerca americana* | 23 | 23 | (Palacios-Gimenez et al. 2020) |
| Acrididae | *Schistocerca damnifica* | 23 | 23 | (Palacios-Gimenez et al. 2020) |
| Acrididae | *Schistocerca ceratiola* | 23 | 23 | (Palacios-Gimenez et al. 2020) |
| Acrididae | *Schistocerca rubiginosa* | 23 | 23 | (Palacios-Gimenez et al. 2020) |
| Acrididae | *Allotruxalis strigata* | 23 | 23 | (Mesa et al. 1982) |
| Acrididae | *Cocytolettix argentina* | 23 | 23 | (Mesa et al. 1982) |
| Acrididae | *Hyalopteryx rufipennis* | 23 | 23 | (Mesa et al. 1982) |
| Acrididae | *Metaleptea brevicornis adspersa* | 23 | 23 | (Mesa et al. 1982) |
| Acrididae | *Parorphula gramínea* | 23 | 23 | (Mesa et al. 1982) |
| Acrididae | *Trimerotropis ochraceipennis* | 23 | 29 | (Mesa et al. 1982) |
| Acrididae | *Trimerotropis pallidipennis* | 23 | 27 | (Mesa et al. 1982) |
| Acrididae | *Amblytropidia australis* | 23 | 23 | (Mesa et al. 1982) |
| Acrididae | *Apolobamba n. sp. prope pulchra* | 23 | 23 | (Mesa et al. 1982) |
| Acrididae | *Dichromorpha australis* | 23 | 23 | (Mesa et al. 1982) |
| Acrididae | *Dichromorpha australis* | 23 | 23 | (Mesa et al. 1982) |
| Acrididae | *Euplectrotettix sp. Nº2* | 23 | 23 | (Mesa et al. 1982) |
| Acrididae | *Euplectrotettix sp. Nº3* | 23 | 23 | (Mesa et al. 1982) |
| Acrididae | *Fenestra bohlsii* | 23 | 23 | (Mesa et al. 1982) |
| Acrididae | *Isonyx paraguayensis* | 23 | 23 | (Mesa et al. 1982) |
| Acrididae | *Isonyx n. sp. Nº1* | 23 | 23 | (Mesa et al. 1982) |
| Acrididae | *Laplatacris díspar* | 23 | 23 | (Mesa et al. 1982) |
| Acrididae | *Meloscirtus montanus* | 23 | 23 | (Mesa et al. 1982) |
| Acrididae | *Notopomala glaucipes* | 23 | 23 | (Mesa et al. 1982) |
| Acrididae | *Orphulella concinnula* | 23 | 23 | (Mesa et al. 1982) |

| | | | | |
|---|---|---|---|---|
| Acrididae | *Orphulella punctata* | 23 | 23 | (Mesa et al. 1982) |
| Acrididae | *Orphulella sp* | 23 | 23 | (Mesa et al. 1982) |
| Acrididae | *Orphulina pulchella* | 23 | 23 | (Mesa et al. 1982) |
| Acrididae | *Parapellopedon instabilis* | 23 | 23 | (Mesa et al. 1982) |
| Acrididae | *Pellopedon sp. (prope obscurum Br.)* | 23 | 23 | (Mesa et al. 1982) |
| Acrididae | *Peruvia nigromarginata* | 23 | 23 | (Mesa et al. 1982) |
| Acrididae | *Scyllina humilis* | 23 | 23 | (Mesa et al. 1982) |
| Acrididae | *Scyllina humilis* | 23 | 23 | (Mesa et al. 1982) |
| Acrididae | *Scyllina signatipennis* | 22 | 23 | (Mesa et al. 1982) |
| Acrididae | *Scyllinops brunneri* | 23 | 23 | (Mesa et al. 1982) |
| Acrididae | *Scyllinops pallida* | 23 | 23 | (Mesa et al. 1982) |
| Acrididae | *Scyllinops n. sp. Nº1* | 23 | 23 | (Mesa et al. 1982) |
| Acrididae | *Scyllinops n. sp. Nº2* | 23 | 23 | (Mesa et al. 1982) |
| Acrididae | *Silvitettix concolor* | 23 | 23 | (Mesa et al. 1982) |
| Acrididae | *Sinipta acuta* | 23 | 23 | (Mesa et al. 1982) |
| Acrididae | *Sinipta acuta* | 23 | 23 | (Mesa et al. 1982) |
| Acrididae | *Sinipta dalmani* | 23 | 23 | (Mesa et al. 1982) |
| Acrididae | *Sinipta maldonadoi* | 23 | 23 | (Mesa et al. 1982) |
| Acrididae | *Staurorhectus longicornis* | 23 | 23 | (Mesa et al. 1982) |
| Acrididae | *Phaulacridium vittatum* | 23 | 23 | (Webb and Westerman 1978) |
| Acrididae | *Paulinia acuminata* | 23 | 23 | (Mesa et al. 1982) |
| Acrididae | *Marellia remipes* | 23 | 23 | (Mesa et al. 1982) |
| Romaleidae | *Agriacris jucunda* | 23 | 23 | (Mesa et al. 2004) |
| Romaleidae | *Diponthus communis* | 22 | 23 | (Castillo et al. 2010) |
| Romaleidae | *Staleochlora frushtorferi* | 23 | 23 | (Mesa et al. 2004) |
| Romaleidae | *Xyleus laevipes* | 22 | 23 | (Castillo et al. 2010) |
| Romaleidae | *Zoniopoda iheringi* | 22 | 23 | (Castillo et al. 2010) |
| Romaleidae | *Alcamenes clarazianus* | 23 | 23 | (Mesa et al. 1982) |
| Romaleidae | *Antandrus viridis* | 23 | 23 | (Mesa et al. 1982) |

| | | | | |
|---|---|---|---|---|
| Romaleidae | *Chariacris miniacea* | 23 | 23 | (Mesa et al. 1982) |
| Romaleidae | *Chromacris speciosa* | 23 | 23 | (Mesa et al. 1982) |
| Romaleidae | *Chromacris miles* | 23 | 24 | (Mesa et al. 1982) |
| Romaleidae | *Chromacris peruviana* | 23 | 24 | (Mesa et al. 1982) |
| Romaleidae | *Coryacris angustipennis* | 23 | 24 | (Mesa et al. 1982) |
| Romaleidae | *Diponthus argentinus* | 23 | 24 | (Mesa et al. 1982) |
| Romaleidae | *Diponthus clarazianus* | 23 | 24 | (Mesa et al. 1982) |
| Romaleidae | *Diponthus sp* | 23 | 23 | (Mesa et al. 1982) |
| Romaleidae | *Diponthus dispar* | 21 | 23 | (Mesa et al. 1982) |
| Romaleidae | *Diponthus electus* | 21 | 23 | (Mesa et al. 1982) |
| Romaleidae | *Diponthus maculiferus* | 21 | 23 | (Mesa et al. 1982) |
| Romaleidae | *Elaeochlora basalis* | 23 | 23 | (Mesa et al. 1982) |
| Romaleidae | *Elaeochlora brachyptera* | 23 | 23 | (Mesa et al. 1982) |
| Romaleidae | *Elaeochlora trilineata* | 23 | 23 | (Mesa et al. 1982) |
| Romaleidae | *Elaeochlora viridicata* | 23 | 23 | (Mesa et al. 1982) |
| Romaleidae | *Elaeochlora sp* | 23 | 23 | (Mesa et al. 1982) |
| Romaleidae | *Eutropidacris collares* | 23 | 23 | (Mesa et al. 1982) |
| Romaleidae | *Prionolopha serrata* | 23 | 23 | (Mesa et al. 1982) |
| Romaleidae | *Procolpia minor* | 23 | 23 | (Mesa et al. 1982) |
| Romaleidae | *Securigera acutangula* | 23 | 23 | (Mesa et al. 1982) |
| Romaleidae | *Xestotrachelus robustus* | 23 | 23 | (Mesa et al. 1982) |
| Romaleidae | *Xyleus attenuatus* | 23 | 23 | (Mesa et al. 1982) |
| Romaleidae | *Xyleus discoideus* | 23 | 23 | (Mesa et al. 1982) |
| Romaleidae | *Xyleus gracilis* | 23 | 23 | (Mesa et al. 1982) |
| Romaleidae | *Xyleus insignis* | 23 | 23 | (Mesa et al. 1982) |
| Romaleidae | *Xyleus modestus* | 23 | 23 | (Mesa et al. 1982) |
| Romaleidae | *Zoniopoda hempeli* | 23 | 23 | (Mesa et al. 1982) |
| Romaleidae | *Zoniopoda juncorum* | 23 | 23 | (Mesa et al. 1982) |
| Romaleidae | *Zoniopoda omnicolor* | 23 | 23 | (Mesa et al. 1982) |

| | | | | |
|---|---|---|---|---|
| Romaleidae | *Zoniopoda similis* | 23 | 23 | (Mesa et al. 1982) |
| Romaleidae | *Zoniopoda tarsata* | 23 | 23 | (Mesa et al. 1982) |
| Ommexechidae | *Neuqeunina fictor* | 22 | 25 | (Castillo et al. 2010) |
| Ommexechidae | *Pachyosa signata* | 22 | 23 | (Castillo et al. 2010) |
| Ommexechidae | *Spathalium audouini* | 22 | 26 | (Castillo et al. 2010) |
| Ommexechidae | *Spathalium helios* | 22 | 25 | (Castillo et al. 2010) |
| Ommexechidae | *Tetrixocephalus willemsei* | 22 | 25 | (Castillo et al. 2010) |
| Ommexechidae | *Tetrixocephalus chilensi* | 23 | 25 | (Mesa et al. 1982) |
| Ommexechidae | *Tetrixocephalus micropterus* | 23 | 25 | (Mesa et al. 1982) |
| Ommexechidae | *Tetrixocephalus sergioi* | 23 | 25 | (Mesa et al. 1982) |
| Ommexechidae | *Tetrixocephalus sp* | 23 | 25 | (Mesa et al. 1982) |
| Ommexechidae | *Aucacris bullocki* | 23 | 25 | (Mesa et al. 1982) |
| Ommexechidae | *Calcitrena maculosa* | 23 | 25 | (Mesa et al. 1982) |
| Ommexechidae | *Clarazella maculosa* | 23 | 25 | (Santander et al. 2021) |
| Ommexechidae | *Clarazella bimaculata* | 23 | 23 | (Santander et al. 2021) |
| Ommexechidae | *Clarazella patagoba* | 23 | 25 | (Mesa et al. 1982) |
| Ommexechidae | *Conometopus sulcaticollis* | 25 | 25 | (Mesa et al. 1982) |
| Ommexechidae | *Cumainocloidus cordillerae* | 23 | 25 | (Mesa et al. 1982) |
| Ommexechidae | *Descampsacris serrulata* | 23 | 25 | (Mesa et al. 1982) |
| Ommexechidae | *Graea horrida* | 23 | 25 | (Mesa et al. 1982) |
| Ommexechidae | *Ommexecha ?* | 23 | 29 | (Mesa et al. 1982) |
| Ommexechidae | *Ommexecha virens* | 23 | 25 | (Santander et al. 2021) |
| Ommexechidae | *Ommexecha macropterum* | 23 | 25 | (Santander et al. 2021) |
